# Capturing the mechanosensitivity of cell proliferation in models of epithelium

**DOI:** 10.1101/2023.01.31.526438

**Authors:** Kevin Höllring, Lovro Nuić, Luka Rogić, Sara Kaliman, Simone Gehrer, Carina Wollnik, Florian Rehfeldt, Maxime Hubert, Ana-Sunčana Smith

## Abstract

Despite the primary role of cell proliferation in tissue development and homeostatic maintenance, the interplay between cell density, cell mechanoresponse, and cell growth and division is not yet understood. In this article we address this issue by reporting on an experimental investigation of cell proliferation on all time- and length-scales of the development of a model tissue, grown on collagen-coated glass or deformable substrates. Through extensive data analysis, we demonstrate the relation between mechanoresponse and probability for cell division, as a function of the local cell density. Motivated by these results, we construct a minimal model of cell proliferation that can recover the data. By parametrizing the growth and the dividing phases of the cell cycle, and introducing such a proliferation model in dissipative particle dynamics simulations, we recover the mechanoresponsive, time-dependent density profiles in 2D tissues growing to macroscopic scales. The importance of separating the cell population into growing and dividing cells, each characterized by a particular time scale, is further emphasized by calculations of density profiles based on adapted Fisher-Kolmogorov equations. Together, these results show that the mechanoresponse on the level of a constitutive cell and its proliferation results in a matrix-sensitive active pressure. The latter evokes massive cooperative displacement of cells in the invading tissue and is a key factor for developing large-scale structures in the steady state.

---

Cell proliferation is the process by which the number of cells in a tissue increases, and is thus implicated in tissue growth, regeneration and homeostasis. The proliferation process comprises the cell growth in size and the cell division into daughter cells, when the cell DNA is duplicated (1). The division itself is tightly controlled by biochemical signaling pathways (2, 3), and has not been found sensitive to details of the cellular environment (4, 5). The regulation of the growth phase has, however, proven to be more delicate and sensitive to mechanical stresses arising from cell-cell and cell-matrix interactions (6–8), quantified through measurements of cell size (9), traction forces, (10–12) and through response to external stresses (9, 13, 14). The emergent conclusion is that increased proliferation depends on a shorter growing phase, while smaller cells experience stronger pressure, delaying their entrance into the division part of the cell cycle (7, 15).

It remains, however, unclear whether in confluent tissues the stiffness of the matrix has a direct impact on the cell’s growth phase, or whether the effects of the matrix are integrated into local stresses from neighboring cells, therefore indirectly affecting proliferation. A logical progression from this question is an inquiry on the contingency of cell proliferation on the developmental stage of the tissue and the position of cells within the emergent tissue compartments, including the homeostatic state. The latter were already found responsive to the mechanical conditioning during the tissue development (16, 17). However, the structuring of the tissue has not yet been related to mechanosensitive properties of single cells and cells in confluent environments, including cell proliferation. Here we address these issues in a joint experimental and theoretical study by investigating the relation between cell proliferation and development of a simple model epithelium, grown on gels and on glass.

## Influence of the local environment on proliferation in MDCK-II tissues

In order to characterize the probability for proliferation in different mechanical environments, we grow unconstrained epithelium from Madin-Darby Canine Kidney II (MDCK-II) cells on collagen I coated glass or 11 kPa PA gel (see SI section S1, and ref.(16) for details). Confluent circular monolayers typically form 6 hours after seeding, when a characteristic density profile forms at the moving edge of the tissue that starts to radially invade the cell-free part of the substrate. In the growing central compartment the tissue densifies until the homeostatic state is reached typically 4 days after seeding (*ρ*_*h*_ = 6860 cells*/*mm^2^ for glass (resp. 7280 cells*/*mm^2^ for 11 kPa gels)) (16, 18). To determine the fraction of dividing cells in maturing tissues, colonies were stained with EdU to highlight the S-phase of the cell cycle (fig. 1a) on day 2, 3, 4 and 6 after seeding. The same colonies are then fixed and stained with Hoechst dye to determine the average cell density profiles as well as local cell densities with high accuracy following previously established protocols (16, 19). The entire fixed tissues are imaged in epifluorescence using 5× and 20× objectives on a Zeiss Cell Observer Z1 microscope.

**Fig. 1.**
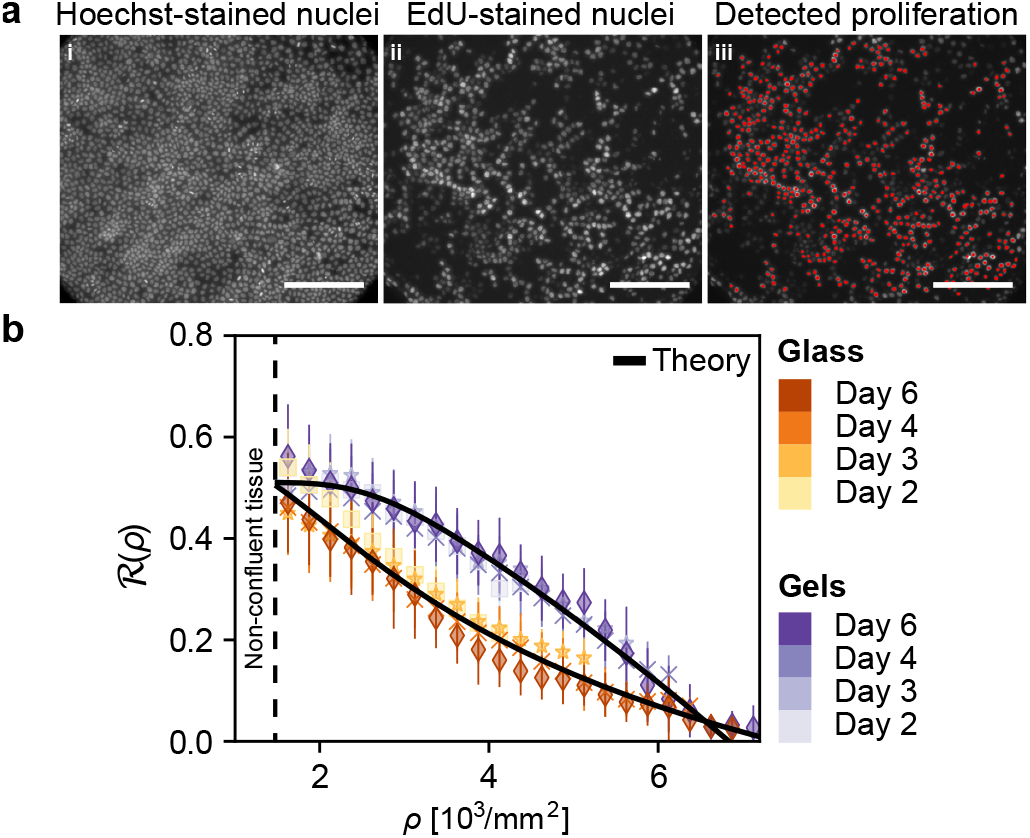
Division of MDCK-II cells in confluent tissues. (a) Images of cells grown on 11 kPa gels stained with Hoechst (i) and with EdU (ii) after 4 days of unperturbed development. A home-made MATLAB post-treatment software highlights the dividing cells based on their EdU-intensity relative to the background (iii, SI section S1). Scale bar = 200 μm (b) Fraction of dividing cells ℛ as a function of the cell density for tissues grown on glass (orange) and 11 kPa gels (purple) Different sampling time points are presented by different symbols. The solid lines are fits of the microscopic model, with the explicit fitting parameters summarized in table S1.

We determine the proliferation probability as the ratioℛ of dividing cells relative to the total number of cells within a region of interest. The size of the region of interest is chosen to be 271 × 241 μm^2^, which accounts for the geometry of the sampled microscopy image and is of the order of the density-density correlation length for MDCK cells (20). The proliferation probability is mathematically defined as

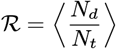

where *N*_*d*_ refers to the number of cells with a positive EdU staining, while *N*_*t*_ is the total number of cells as determined from Hoechst staining (fig. 1a). The brackets ⟨· ⟩ denote the ensemble average performed over the bin. In total, we sample 10^4^ regions of interest (see fig. S1), which were consequently binned in cell density windows of 250 cells*/*mm^2^ (fig. 1b).

Inspection of fig. 1b shows that the fraction of dividing cells decreases in a non-linear fashion as the local cell density increases. Ultimately, ℛ goes down to almost zero when the homeostatic density is reached on days 4 and 6, as found previously (9, 15). On gels, there are basically no proliferating cells in the homeostasis, while on glass, some residual divisions occur, due to the larger amount of extrusions (16).

Another important finding in fig. 1b is that at fixed substrate stiffness, data for all days overlap (see also fig. S2). This suggests that ℛ is only a function of the density in the immediate environment of the dividing cell, and is not sensitive to the history and spatial positioning of the cell within the tissue. The time merely sets the accessible density range (fig. S2). The origin of this behavior is likely related to YAP signaling (8), which was recently found to be more localized in the nuclei under lower confinement, whereas at high confinement, it moves to the cytoplasm (21).

The data also clearly shows that cell proliferation explicitly depends on the stiffness of the underlying substrate. Notably, the probability of cell division in tissues grown on glass displays a curve with a positive (convex) curvature, while in tissues grown on gels, ℛ has a negative (concave) curvature (fig. 1b), which indicates more probable cell proliferation on gels than on glass at all densities. These mechanosensitive features clearly implicate integrin-mediated adhesions in the regulation of proliferation, which likely emerges from the smaller tension that integrin adhesion generates on soft substrate than on hard substrates. This is in turn compensated by E-cadherin intracellular binding and the formation of apical actin structures, which are more intensive on soft substrates than on hard ones (13, 22). This obviously affects the homeostatic state of the tissue (16, 23), and as seen here, the growth phase of cells.

### Microscopic model for cell proliferation

To rationalize these experimental findings, we devise a minimal cell-level model for proliferation. Following previously suggested strategies (24), we divide the cell cycle into a growth phase and a division phase, each having a characteristic duration, denoted respectively by *τ*_*g*_ and *τ*_*d*_. These time scales need to be determined to estimate the observable ℛ

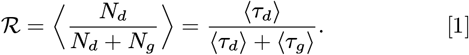

within a region of constant local cell density consisting of *N*_*g*_ growing and *N*_*d*_ dividing cells (*N*_*t*_ = *N*_*g*_ + *N*_*d*_).

We first realize that the time dependent size of a cell in a tissue *σ*(*t*) can only evolve in a range *σ*_0_ ≤ *σ*(*t*) ≤ *σ*_*ρ*_, where *σ*_0_ is the size of a daughter cell and *σ*_*ρ*_ is the maximum size that reflects the local density *ρ* of cells applying an isotropic pressure on the proliferating cell (fig. 2a). As the tissue matures *ρ* increases until the homeostasis is achieved, characterized by the density *ρ*_*h*_ *> ρ* and cell size *σ*_*h*_ (fig. 2b).

**Fig. 2.**
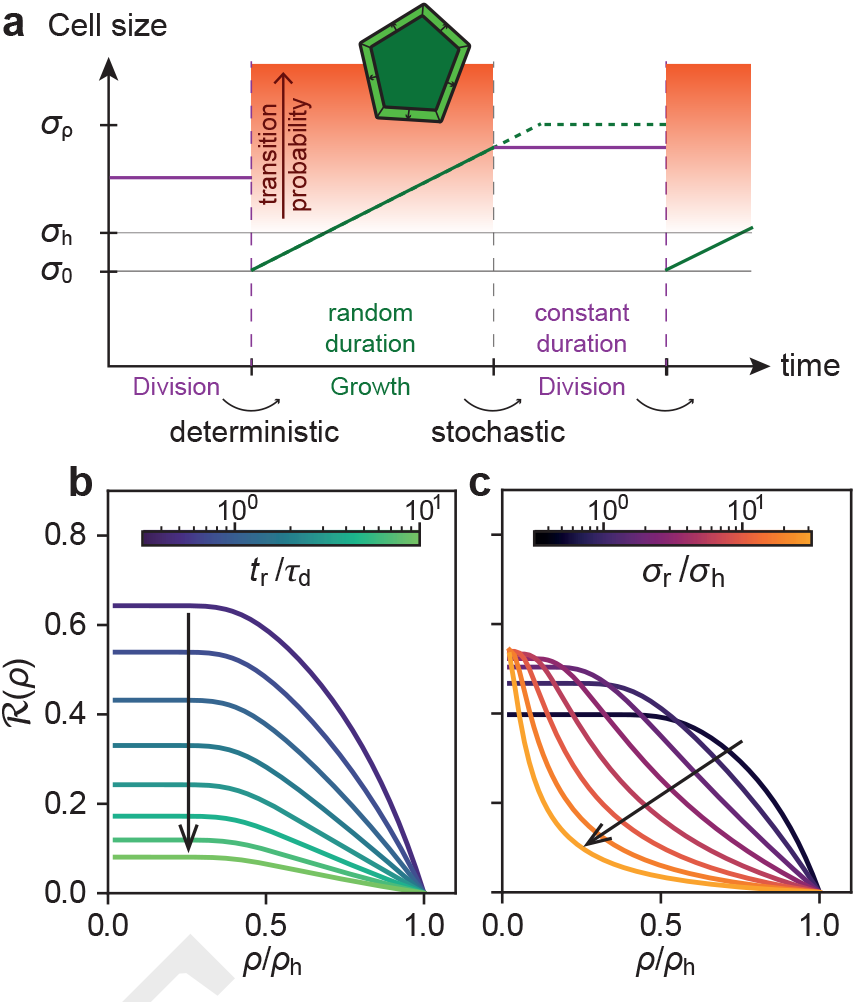
Microscopic model for cell proliferation. (a) The different stages of the cell life cycle in our models. The cell grows in size while in the *growing* stage and transitions stochastically towards the *division* stage. It then remains at constant size for a deterministic amount of time before producing two daughter cells. The graph shows the cell size over time, the duration of each phase and the probability to transition between the growth an division. (b) Impact of changing the the ratio of *t*_*r*_*/τ*_*d*_, which controls the absolute amplitude ℛ. (c) Modulation of *σ*_*r*_ on ℛ shows that this parameter mainly controls the curvature of ℛ through the point of inflection of *τ*_*g*_ (see fig. S3).

We first account for the time scale characterizing the *division phase* ⟨*τ*_*d*_⟩. Following reports in the literature about its robustness in different environments (4, 5, 21), we set ⟨*τ*_*d*_⟩ to be constant. Furthermore, we assume that a cell’s transition to the growth phase takes place in a deterministic manner after ⟨*τ*_*d*_⟩ in all density backgrounds.

To model the *mechanosensitive growth phase* we assume a linear growth law for the time evolution of the cell size *σ*(*t*)

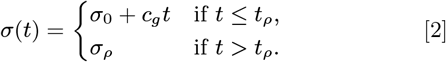

Here, *c*_*g*_ is a constant and defines the speed at which the cell grows and *t*_*ρ*_ is the time at which the cell attains *σ*_*ρ*_. In calculations, the local density background *ρ* is kept homogeneous such that only the cell of interest is able to grow as a function of the time *t*. This is justified both at low and high densities. For *ρ* ≪*ρ*_*h*_, one has *τ*_*d*_ ∼ *τ*_*g*_ and there are only a few cells actively growing. For *ρ* ≃*ρ*_*h*_, *τ*_*d*_ ≪*τ*_*g*_, and most cells are in their growth phase, but their total change in size is actually small.

To model the transition from the growth to the division phase, we first discuss the rate *r*_*d*_(*σ*) at which cells stop growing and enter the division phase in a stochastic manner (25). To capture the effect of pressure induced by neighbors in the confluent tissue, we model *r*_*d*_(*σ*) as a simple function of the cell size:

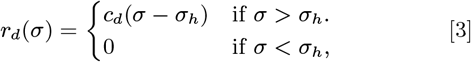

The constant *c*_*d*_ defines the rate with which *r*_*d*_ increases with *σ*(*t*). If the cell is much larger than a cell in the homeostatic state (*σ*− *σ*_*h*_) ≫ 0, the rate of division is high. On the other hand, if *σ*(*t*) → *σ*_*h*_, then *r*_*d*_ → 0. As such, a vast majority of cells starts to divide before attaining their maximum possible size. To calculate the average time that the cell spends in the growth phase ⟨*τ*_*g*_⟩, we determine the change in the probability *p*_*g*_ for a cell to still be in the growing phase at time *t >* 0. This change is simply a product of the probability that the cell is still growing (*p*_*g*_) and the rate of division associated with the attained size:

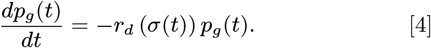

The average time ⟨*τ*_*g*_⟩ spent growing can thus be calculated by combining eq. (4) with eqs. (2) and (3), which yields

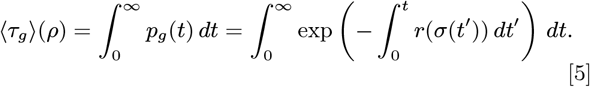

Following the calculation shown in SI section S2, we find

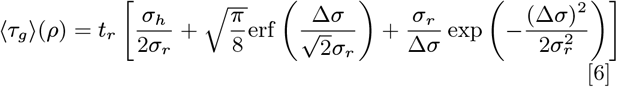

with Δ*σ*= *σ*_*ρ*_ − *σ*_*h*_ bringing the explicit tissue-density dependence of ⟨*τ*_*g*_ ⟩ through *σ*_*ρ*_ (see SI section S2 for details).

The parameters 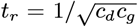 and 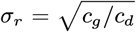, are the characteristic time and length scales for the mechanoresponse of the growth phase and in essence encode the mechanosensitivity of individual cells. They also determine the degree to which variations in tissue pressure constrain the cell growth, thereby linking single cell properties to properties of cells in a tissue environment, as suggested in earlier works (26). Consequently, larger *σ*_*r*_ comes with larger cell sizes at time of division and can also be linked to larger cell sizes in homeostasis in our experiments. More specifically, it also controls the predicted excess size of an isolated cell at time of division relative to the cell size in homeostasis *σ*_*h*_ (see SI section S3).

The determined ⟨*τ*_*g*_⟩ is found to increase monotonously with tissue density (see fig. S3)in a non-linear fashion. This behavior is consistent with our experiments and recent density dependent measurements of *τ*_*g*_ (9, 21). Close to *ρ*_*h*_, ⟨*τ*_*g*_⟩ rises dramatically, corresponding to the contact inhibition of proliferation in homeostasis (8).

#### Comparison with experiments

With our model for ⟨*τ*_*g*_⟩ and the considerations for ⟨*τ*_*d*_⟩, we can now derive the fraction of dividing cells using ℛ eq. (1) (fig. 2b and fig. 2c). The scale *σ*_*r*_ controls the curvature through the point of inflection of ℛ (*ρ*) (fig. 2c), while the amplitude of the curve towards lower densities is mainly tied to *t*_*r*_ (fig. 2b; see section S2 for details).

Our calculated ℛ only relies on the relative scale of the duration of the two phases and not on the respective absolute time, hence the precise shape of ℛ (*ρ*) only depends on the parameters *t*_*r*_*/* ⟨*τ*_*d*_⟩, *σ*_*r*_ and *ρ*_*h*_. Therefore, fitting of experiments (fig. 1(b)) with eq. (1) requires only two parameters along with the homeostatic density. The latter, *ρ*_*h*_, can be extracted from the fit (which yields 6800 cells*/*mm^2^ to 7400 cells*/*mm^2^), or it can be fixed to the values that have been measured with high precision (6860(360) and 7280(260) cells*/*mm^2^ for glass and 11 kPa gels, see fig. S4) (16).

The fit results (see SI table S1 and section S4) provide a prediction for the area of the lone cell at time of division ⟨*σ*⟩ of 780.1 μm^2^ on glass and 374.0 μm^2^ on gels. This is in very good agreement with measured 2000(900) μm^2^ on glass (27) and a 30 % to 40 % decrease for cells on gels (28).

It is natural to expect that cells in the tissue on gels are generally smaller than those on glass based on the smaller value of *σ*_*r*_ obtained from our fit, which is indeed the case, both in the homeostatic state and in the edge compartments (16). This result is further corroborated by the analysis of rapidly spreading MCF-7 at the single cell level when grown on PAAm gels for a fixed duration of 12 h (29).

The fit furthermore concludes that the ratio *t*_*r*_*/*⟨*τ*_*d*_⟩ on glass takes the value of 0.91 and 0.90 on gel. The appropriateness of the first result can be validated through the use of experimental results on glass substrates of a full cell cycle time *τ*_*g*_ + *τ*_*d*_ of 15 h (see SI fig. S5) and the recently reported duration of cell division (S, G_2_ and M phase) in similar conditions of about 10 hours (21). As detailed in SI section S4, this yields an experimental estimate of *t*_*r*_*/τ*_*d*_ = 0.83 which is in very reasonable agreement with the fit itself.

By assuming that only the growing phase is mechanoresponsive, and that *τ*_*d*_ is insensitive to the mechanical environment (21), our model predicts a longer growth time *τ*_*g*_ on glass than on gels (see SI section S4). In experiments, despite the fact that the cells within a tissue are indeed smaller on gel substrates than on glass, we also observe that the colonies on gels are larger. Consequently the total number of cells on gels after 6 days is larger even though the seeding conditions are identical. This implies an actually shorter cell cycle, as predicted by our model.

Notably, simpler division protocols, such as a deterministic cell division above a given threshold (30–35) or a constant probability to start division above a given threshold (36), do not reproduce the features of the experiments in fig. 1b (see SI section S5, figs. S6 and S7). The here proposed model has, however, the capacity to fit the data and recover reasonable values for both *σ*_*r*_ and *t*_*r*_*/τ*_*d*_. The model also shows that the mechanoresponse of cells in a tissue may be directly related to the mechanosensitivity of a single cell captured in *σ*_*r*_ and *t*_*r*_.

### Capturing the mechanoresponse of proliferation in DPD simu-lations

As an additional means of verification of our microscopic model, we use Dissipative Particle Dynamics (DPD) simulations (see SI section S6 for details) (30, 31, 33–35, 37– 40). This method, with an instantaneous division phase and a stochastic growth phase duration (30–33), successfully demonstrated the effect of isotropic pressure on spheroids. These simulations showed that externally applied pressure limits cell proliferation, by restraining and even preventing cell growth (30, 31). In order to capture mechanoresponsive properties presented in (fig. 1), the proliferation rule must be modified. Consistently with our minimal model we introduce the two phases of the cell cycle (for formal mapping see SI section S7). For the growth phase, we keep the usual approach for DPD simulations of tissues. Specifically, an inner cell force *F*_*g*_ = *B/*(*l* + *r*^⋆^)^2^ is defined to model the cell growth by increasing the distance *l* between the particles representing the cell (fig. 3a). Here, *B* is the force intensity and *r*^⋆^ an offset to avoid numerical divergence. We, furthermore, introduce a *division* phase, where we cancel *F*_*g*_ and instead introduce a force *F*_*d*_ = *K*_*d*_(*l* − *l*^⋆^) to fix the size of the cell during division (fig. 3a). Here, *K*_*d*_ is the cell stiffness and *l*^*⋆*^ is the cell size when entering the division phase. The cell remains in the division phase for a constant duration *τ*_*d*_. After this time expires, two daughter cells are produced with an initial particles distance *l* = *l*_0_, re-initiating the growth phase. Following the ingredients used in the microscopic model, we define the probability to enter the division phase as

**Fig. 3.**
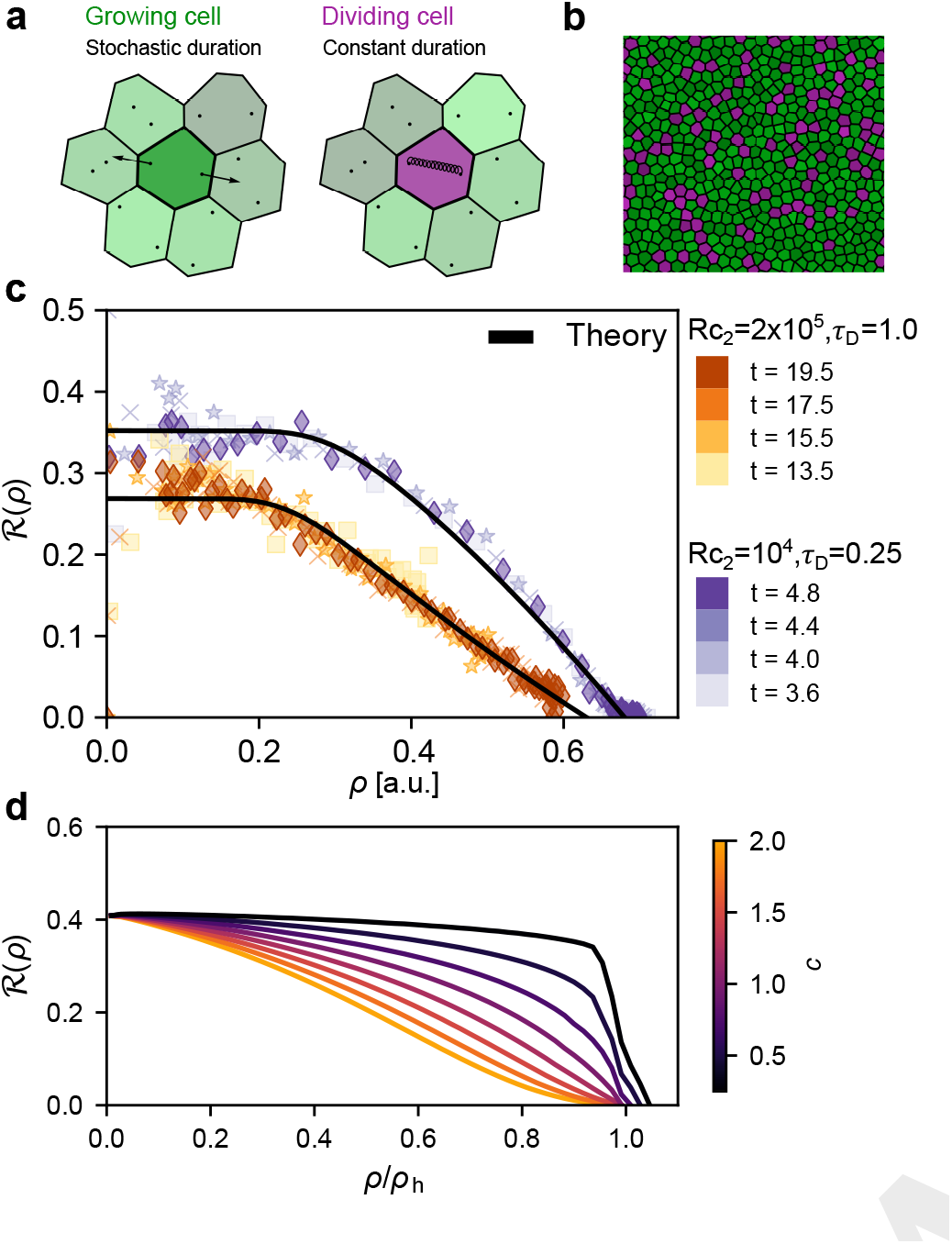
Quantification of proliferation in numerical simulations of expanding epithelia. (a) Schematics illustrating the inner forces within cells in the DPD simulations depending on their state, i.e. *growing* or *dividing*. (b) Snapshot illustrating the dividing cells within the expanding tissue. *Growing* cells are represented with shades of green while *dividing* cells are represented with shades of purple. (c) Fraction of dividing cells in the tissue as a function of the local density for two simulations with different values of *R*_*c*2_ and *τ*_*d*_ as indicated in the legend. Different instants are considered as highlighted by the different shades of green and pink. All other parameters used in the simulations are indicated in Ref. (16). (d) Dependence of ℛ (*ρ*) on the choice of *c* in the DFK formalism. As desired, *c* changes the curvature of the graph towards the homeostatic density *ρ*_*h*_. We also observe slight changes in the observed convergence density, which deviates slightly from *ρ*_*h*_ consistent with our experimental observations of density bumps on soft substrates.

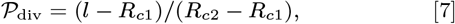

where *R*_*c*1_ and *R*_*c*2_ are two size thresholds, and changing their *R*_*c*2_ − *R*_*c*1_ corresponding to changes in *σ*_*r*_. This probability is evaluated at each time step of the simulation for each cell in the growth phase.

Mimicking the previously-described experiments, we perform simulations of freely expanding monolayers in two dimensions. Thereby, a small patch of a tissue is placed in the center of the simulation box, with cells being at low density. With time, the tissue increases its density and expands, developing a density profile. Ultimately, the central compartment reaches homeostasis. Furthermore, to affect the ratio *t*_*r*_*/τ*_*d*_, we vary *τ*_*d*_, instead of *t*_*r*_, as the former is explicitly defined in DPD, whereas the latter is only implicit. To be consistent with experiments, we periodically measure ℛ, and sample the fraction of dividing cells within regions of constant cell density, throughout the tissue and at various time steps. We fit eq. (1) to the simulation results (SI table S2), and similarly as in the experiments we see that the microscopic model of cell proliferation captures precisely the DPD results (fig. 3c). We find that changing *R*_*c*2_ − *R*_*c*1_ and *τ*_*d*_ (to effectively modify *t*_*r*_), as in experiments, leads to an overall curvature change of ℛ, the latter decreasing monotonously from a constant value at low densities towards zero in the homeostatic state (fig. 3c). As expected we find that increasing *R*_*c*2_ increases *σ*_*r*_, which in conjunction with *t*_*r*_ larger than *τ*_*d*_ provides dynamics like in tissues grown on glass, and larger cells in homeostasis. Smaller *R*_*c*2_, with *t*_*g*_ comparable to *τ*_*d*_, provides results comparable to tissues on gels and smaller cells in the homeostatic state. Finally, the resulting ℛ(*ρ*) is independent of both the sampling time and the position within the tissue, as demonstrated by the fact that all curves taken at different snapshots fall onto a common master curve, consistent with the experimental findings (fig. 1). Such results align with our initial assumption about the nature of the transitions between the growth and division phases, which is the key ingredient for the observed behavior.

### Mechanoresponse of the tissue proliferation in the delayed Fisher-Kolmogorov formalism

To verify the robustness of these results, and the role of the non-linearity of proliferation probability, we furthermore develop a theoretical model for tissue growth that incorporates the key ingredients identified here-above. We start from the Fisher-Kolmogorov (FK) theory (41). The evolution of the density profile is captured by an (active) diffusion-like process with efficiency *D*. The cell proliferation is described through a logistic-like growth process with efficiency *β* (41). This model, however, provides only a linear dependence of the fraction of dividing cells on the density. One way to incorporate the nonlinear effects is by introducing complex empirically fitted functions as suggested by Metzner *et al*. (42) for HT1080 fibrosarcoma cells and MCF7 mammary gland adeno-scarcinoma cells. We choose a different strategy to capture mechanosensitivity of proliferation. Instead of modeling the evolution of the *total* cell density, we first distinguish populations of cells in the growth and division phase (*ρ*_*g*_ and *ρ*_*d*_ respectively).

We, furthermore, introduce a non-linearity through an exponent *c* in the logistic term (see SI section S8 for motivation of this form). This parameter is used to control the curvature of ℛ(*ρ*) through a single parameter and thus provides the simplest inclusion of the mechanoresponsiveness (see fig. 3d, SI section S8). We, furthermore, introduce a delay *τ* = *t* − *τ*_*d*_, to account for the time that cells deterministically spend in the division phase. With these considerations we arrive at

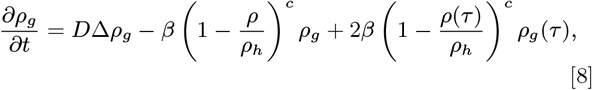

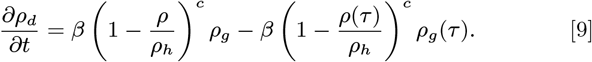

Here, the total density *ρ* = *ρ*_*g*_ + *ρ*_*d*_ is given by the sum of the two above equations leading to

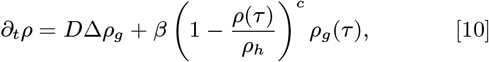

which we term the Delayed Fisher-Kolmogorov (DFK) equation. For the case *c* = 1 and *τ*_*d*_ = 0, eq. (10) amounts to the usual FK model (37).

The parameters of the microscopic model (i.e. *t*_*r*_, *σ*_*r*_, *τ*_*d*_ and *σ*_*h*_) are captured by (*β* ∝ 1*/t*_*r*_, *c, τ*_*d*_ and *ρ*_*h*_ = 1*/σ*_*h*_) (see figs. S8 and S9 for detailed discussion). The set of eqs. (8) to (10) is then solved using a home-made delayed differential equation solver described in the SI section S9.

We first explore the dependence of the fraction of the dividing cells ℛ (*ρ*) = *ρ*_*d*_*/*(*ρ*_*d*_ +*ρ*_*g*_) on the parameter *c*. We find ℛ independent of the position and the age of the colony, and the proliferation probability is again only a function of density. For *c* = 2 we find ℛ similar to the one observed on glass, while for *c* = 0.5 we obtain a dependence resembling the behavior of cells on gels (fig. 3d). Obviously, this one-parameter approach is sufficient to capture the non-linearity (and the curvature) as main feature of ℛ(*ρ*) observed in experiments. Its powerlaw behavior close to the homeostatic density, controlled by *c*, can therefore be seen as a first-order approximation of our microscopic model that suggests a polynomial (see SI section S10).

### Mecanosensitivity of proliferation and the macroscopic organisation of the tissue

We now look at the consequences of nonlinearities in the density dependent proliferation rates, such as shown in fig. 1b, fig. 3c, or fig. 3d, and hypothesize that different mechanosensitivity of proliferation will yield different macroscopic tissue organisation (43). Indeed, the expansion and maturation of tissues on glass and on gels over 6 days show distinctly different outcomes (fig. 4(a,b)), a result that is captured both in simulations (fig. 4(c,d)) and by the DFK formalism (fig. 4(e,f)).

**Fig. 4.**
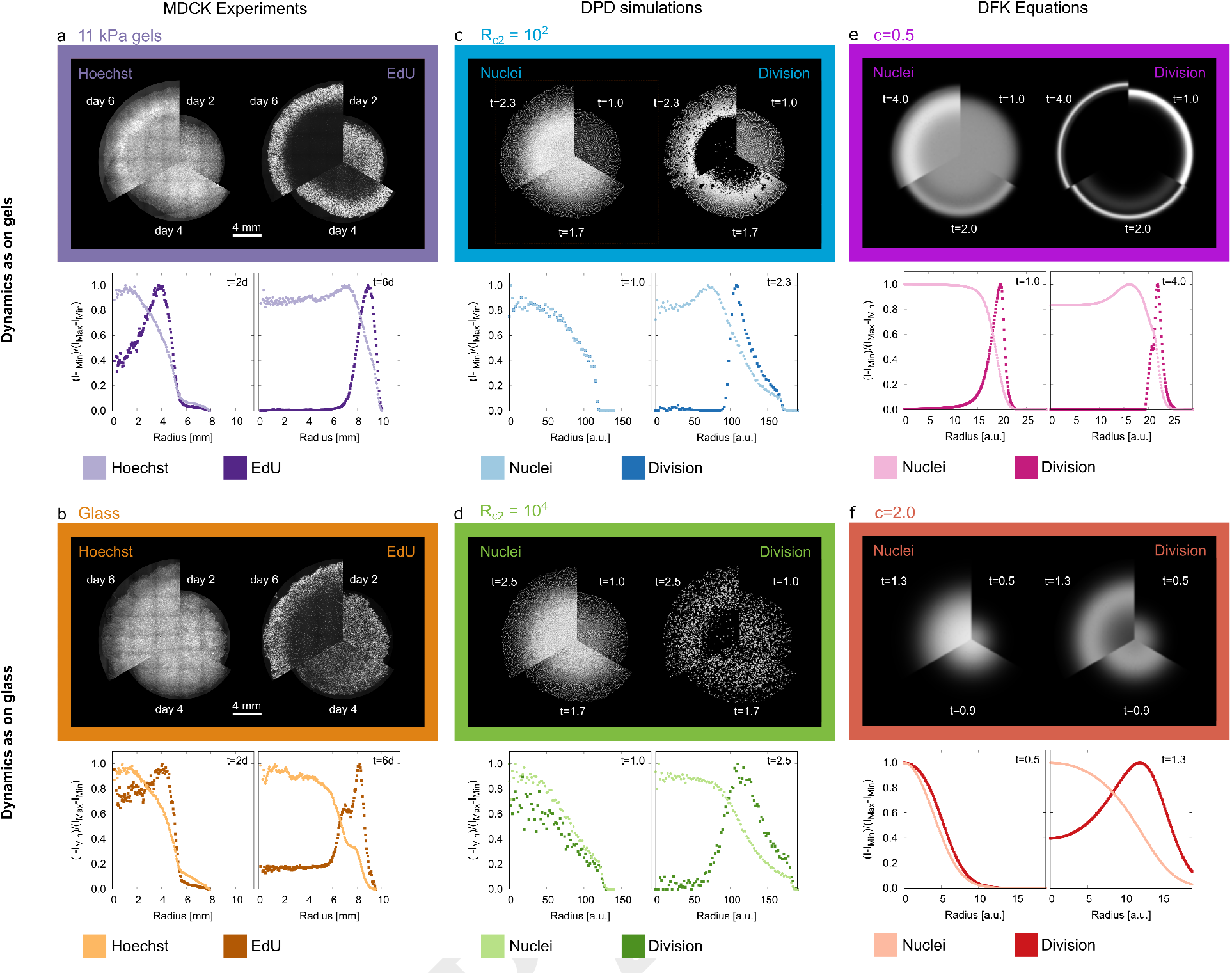
Relation between the evolving macroscopic structure of expanding tissues and the mechanoresponsive proliferation. Panels (a,b) show data from experiments on glass and gels. Panels (c,d) provide equivalent information from DPD simulations, respectively, while (e,f) present the results of the DFK model. Each panel compares the macroscopic tissue structure in early stages of development, close to the formation of the homeostatic state, and after the formation of the steady state expansion profile. For each stage a third of the colony is cut out and juxtaposed with cutouts from other stages. Overall cell distributions (top right) are furthermore compared with the distribution of proliferating cells extracted from the same tissue (top left). The graphs in each panel represent the average radial density of cells and the density of proliferating cells throughout the colony in early (left graph) and steady growth phase (right graph). (a,b) Experimental space-time evolution of the Hoechst and EdU intensities across the tissues. The graphs show normalized radial intensity profiles on day 2 and day 6. (c,d) DPD simulation space-time evolution of cell density (*ρ*_*g*_ + *ρ*_*d*_), and proliferating cell density (*ρ*_*d*_). Mimicking the experimental results is possible by changing the parameters *R*_*c*_2 − *Rc*1 and *τ*_*d*_. (e,f) Equivalent information is obtained from the DFK model, where the experimental results are recovered by changing the parameter *c*. Other parameters are kept constant.

In early stages of development, our model tissues are way below the homeostatic density, and a large concentration of proliferating cells is found throughout the colony. With time, the proliferation becomes less probable in the center where the density of cells increases. Simultaneously, the EdU profile develops a peak which is shifted into the moving edge of the tissue. The peak is the simple consequence of the fact that cells at high density proliferate less than larger cells in low density regions where only few cells contribute to the EdU intensity. The peak therefore denotes the density at which the cell number and their probability to divide is optimal.

Due to the concave shape of ℛ, this peak is more pronounced on gels than on glass, where ℛ is convex, a fact particularly well highlighted in the DFK, due to the lack of statistical fluctuations. Six days after seeding, the homeostatic state appears in the central compartment, which is evident from a strong drop in the EdU signal, but also by the inhibition of locomotion, also observed in DPD and in DFK calculations.

The most notable feature in the tissues growing on gels (but not on glass) is the appearance of an overshoot in the density profile, found in experiments and spontaneously appearing in DPD and DFK models. It appears at the edge of the central homeostatic compartment, and is characterized by a cell density larger than *ρ*_*h*_. Within the overshoot, proliferation is basically inhibited, however, cells are still moving. The robustness of this result suggest that the overshoot is a direct consequence of the finite, deterministic time of cell division, supported by slow density equilibration enabled by active transport. Simply said, cells will begin dividing when they are near the homeostatic state, and they will continue the process even if their doubling causes the density to exceed the homeostatic level. If motility is low, as the measurements suggest, the build-up of cells transiently appears. Due to the shape of ℛ (*ρ*), close to the homeostatic more cells will enter the division state on gels than on glass, leading to the difference in the density profiles, as evidenced by experiments, simulations and our theoretical modeling. These results clearly demonstrate that appropriate modeling of proliferation is crucial for the correct recovery of macroscopic structures during development.

## Discussion and conclusions

From a theoretical point of view, our cell-level model aligns with the description of Smith and Martin of single cell proliferation (25, 44), which is used in the case of immune cells (45), but here our contribution gives a mechanoresponsive twist. Indeed, using eq. (6), one can derive the statistics of time spent in the growth phase (see SI section S2.section C for details), the so-called *α*-curve (44), using Equations (2) to (4). Importantly, these distributions can be related to the parameters of our mechanoresponsive microscopic model, and consequently to the mechanoresponsiveness of individual cells. An independent validation of our model comes from successfully applying the fit function obtained from our data, to results on the same cell line provided by others in the field (21).

Beyond the experiments discussed in this article, our theoretical framework can shed some light on the response of dense tissues to stretching (9, 13), which was found to induce a burst of division events, even reactivating proliferating dynamics despite having been at homeostasis (9). Such results can be easily understood using our microscopic model, where stretching can simply take the cells into the regime where *r*_*d*_(*σ*)≠ 0 (see eq. (4)). Naturally, the opposite experiment, namely compression (30, 31, 46), could lead to an arrest of proliferation by taking the cells to the regime where *r*_*d*_(*σ*) = 0 depending on the compression amplitude. Our microscopic model of proliferation can provide the delay upon which the burst of dividing cells reaches its maximum as a function of the stretching amplitude, as well as the point of arrest due to compression.

Our model is also consistent with results of Abdul Kafi *et al* (47) on HeLa (human cervical) and HEK293T (human embryonic kidney) cells grown on functionalized nano-dot, nano-rods and nano-pillars. They measured increased proliferation for taller structures which was not understood. An increase of proliferation was also observed when changing the diameter of the nanopillars (48). With the length of nano-pillars inversely proportional to their stiffness, our model suggest that the enhanced proliferation actually relates to mechanotransduction of proliferation - taller structures appear as softer substrates, while smaller adhesive areas decrease integrin induced stresses, providing similar effects.

Finally, we believe that our study will shed a new light on the interaction of proliferation and maintenance of homeostasis in more complex epithelia. Indeed, the mechanical properties of the basal membrane are changed for example in various skin diseases which has been related to the proliferation behavior of cells (49, 50). The here suggested models could provide deeper insights into the relation between observed proliferation, and the emerging restructuring of the tissue due to mechanoresponse.

In closing, the main contribution of our work is the demonstration of the dependence of the proliferation probability on the mechanical properties of the microenvironment. The proliferation probability was evaluated in tissues during all stages of growth - from low density seeding to homeostasis. We experimentally demonstrated that the fraction of proliferating cells only relates to the local density. This observed behavior is captured by a minimal microscopic model for mechanosensitive cell proliferation in tissues. The model demonstrates that the shape of the relation between proliferation rate and density on a tissue level can be captured purely trough the description of the explicit mechanoresponse of the proliferation of individual cells and the homeostatic density. The mechanosensitivity of the proliferation probability is then a consequence of cells experiencing local environmental pressure set among others by the varying local density of cells. This result is fully consistent with our experimental findings, which then show that the local proliferation probability in tissues does not depended on the macroscopic state of the tissue during its development, as long as the mechanical environment stays unaltered.

The effect of proliferation on the macroscopic structuring of tissues is further studied by adapting the microscopic model to DPD simulations and capturing its essence in the delayed Fisher-Kolmogorov approach. Besides highlighting the role of a mechanosensitive growth phase and a robust, deterministic division phase in the life cycle of the cell, these approaches, in full agreement with experiments, show that the evolution of density and size of a tissue is indeed deeply affected by mechanoresponsive cell proliferation.

This work opens a new perspective on cell proliferation and on the theoretical description of developing epithelial tissues. In the near future, we can hope this new detailed description will provide a better understanding on the growth and development of more realistic epithelial tissues. Furthermore, we hope that this work will provide a new perspective on epithelia-related diseases and their eventual relations to alterations of the stiffness of the extracellular matrix.

## ACKNOWLEDGMENTS

This work was in part supported by the German Research Foundation projects RTG 1962 (ASS, SG), RTG 2415 (ASS, MH, KH), SFB 755 project B08 (FR), and the ERC StG 337593 MembranesAct (ASS, SK, LN).

