## Supplementary Information (SI) for "Capturing the mechanosensitivity of cell proliferation in models of epithelium"

PNAS
www.pnas.org

### Supplementary Information for

#### Capturing the mechanosensitivity of cell proliferation in models of epithelium

**K. Höllring, L. Nuić, L. Rogić, S. Kaliman, S. Gehrer, C. Wollnik, F. Rehfeldt, M. Hubert and A.-S. Smith**

**Ana-Sunčana Smith**
****

##### This PDF file includes:

Supplementary text
Figs. S1 to S9
Tables S1 to S3
SI References

**S3 Effect of $t_r$ and $\sigma_r$ on growth duration and division statistics of isolated cells** **12**

**S4 Validating the cell-level model's predictions against experimental data** **16**

**S5 Issues with simpler models** **17**
A Deterministic proliferation . . . 17
B Constant proliferation rate . . . 19

**S6 Dissipative Particle Dynamics simulation methods** **22**

**S7 Mapping the DPD and cell-level models** **23**

**S8 Motivation of the Delayed Fisher Kolmogorov Formalism** **26**

**S9 Details of the DDE solver and the numerical solution to the DFK** **27**

**S10 Behavior of $\tau_g$ and $\mathcal{R}$ around $\rho_h$** **31**

#### Supporting Information Text

#### S1. Experimental Methods

**Cell Culture and gels preparation.** The preparation of gels and the culture of cells follow the procedure used in a previous publication of our group (1).

MDCK-II cells were obtained from ECACC, UK (Cat #00062107, RRID:CVCL_0424) and cultured in MEM Earle's medium (#F0325, Biochrom) supplemented with 5% fetal bovine serum (FBS, #F0804, Sigma-Aldrich), 2mM L-glutamine (#G7513, Sigma-Aldrich), and 1% penicillin & streptomycin (#15070-063, Gibco, LifeTechnologies) at 37°C and 5% $CO_2$. Cells were passaged every two or three days before reaching 80% confluence.

Elastic polyacrylamide (PA) gels were prepared as follows. Appropriate mixtures of acrylamide (40% solution, BioRad) and bis-acrylamide (2% solution, BioRad) were polymerized by addition of 0.1%(v/v) N,N,N,N tetramethylethylenediamine (TEMED) and 1%(v/v) ammonium persulfate (APS) for 60 minutes at RT on plasma cleaned glass cover slips (No.1, 25mm ∅, VWR) that were pre-treated with 3-aminoproyltriethoxysiliane (APTES, Sigma-Aldrich) for 15 minutes and incubated with a 0.5% solution of glutaraldehyde in PBS (Sigma Aldrich) for 30 minutes. For quality control reasons, we used the stock solutions not longer than 3 months stored at 4°C and protected from light and ideally prepared as many samples from the same batch as needed.

After polymerization the PA gels were washed extensively with PBS and subsequently coated with Collagen-I (BD Biosciences) at 0.02 mg/mL in a 50 mM HEPES buffer using the bi-functional cross-linker Sulfo-SANPAH (Pierce, Thermo Scientific) activated for 10 minutes with UV light (365 nm). Here, we use the conditions optimized in (2), at which the concentration of collagen on the surface is fully saturated, to avoid deviations in available collagen density, and to make sure that adhesive properties of the surfaces are identical in all cases.

**Imaging.** Tissues were fixed using a 10% solution of formaldehyde (formaldehyde, #47608, Sigma-Aldrich) in PBS for 5 minutes, and the cells were permeabilised for 10 minutes using a 0.5% solution of Triton X 100 (Carl Roth). After washing with PBS, samples were blocked using 3% BSA (#A9418, Sigma-Aldrich) in PBS for 30 minutes at RT, which preceded another Triton X 100 treatment of 5 minutes at RT, followed by 3 washing steps with PBS.

Epifluorescence microscopy images were acquired on an inverted microscope (Zeiss Cell Obsever Z1) using 5× and N-Achroplan 20× objectives, using AxioCam M3 and AxioVision software package (all Zeiss). Confocal microscopy was performed on a Leica LSM SP5 laser-scanning microscope equipped with a white light laser and a 63× and a 100× oil immersion objective yielding fields of view of (246 $\mu$m × 245 $\mu$m) and 100× (155 $\mu$m × 155 $\mu$m) respectively. The step size in the z-direction was kept constant at 0.25 $\mu$m. All tissues grown in one series were fixed and stained simultaneously and imaged consecutively without changing the microscope settings.

**Density and proliferation analysis.** To quantify density and proliferation, we first segmented Hoechst-stained-nuclei picture using MATLAB. From the number of the nuclei center of mass we can evaluate the cell density, and from the position of the center of mass, we can evaluate the cell proliferation using the EdU-stained-nuclei pictures.

Firstly, 5% of the original $3.2 \times 2.7$ mm$^2$-image is removed at the edge because of the overlap between adjacent images in the overall picture of the cluster. Following that step, the reduced images are divided in 108 segments with size $271 \times 241$ $\mu$m$^2$ to avoid segmentation error due to a single thresholding parameter being applied to the whole image. On each of the 108 sub-pictures, the Otsu method is used to determine the optimal thresholding parameter. From the binary images resulting from segmentation, a distance-transform is applied and followed by a standard watershed algorithm. The final result is a segmentation of nuclei which allows to extract their center of mass. The cell density is then evaluated by taking the number of nuclei divided by the area of the reduced picture. In average, such a procedure provides in-between 130 and 480 cells with area varying from 1500 cells/mm$^2$ up to 7000 cells/mm$^2$.

Secondly, knowing the outline of the nuclei from the Hoechst picture, one can evaluate the EdU intensity on the corresponding EdU picture. If the EdU-intensity at the nucleus position is higher than 95% of the average EdU-intensity of the image segment, the cell is tested positive for proliferation. Knowing the number of dividing cells in the image segment and the total number of cells in the corresponding Hoechst image, one has access to the ratio of proliferating cells.

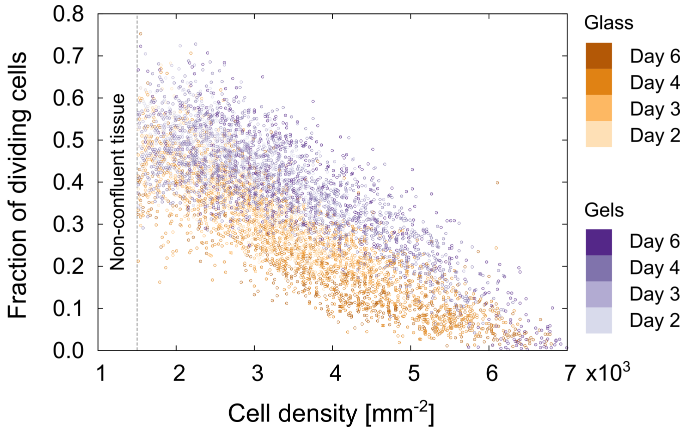

**Fig. S1. Division of MDCK-II cells in confluent tissues and its relation to the cell local environment: Unbinned results.** Presentations of the results shown in Fig. 1 of the main text before binning and averaging. The color used matches the one used in the main text (3).

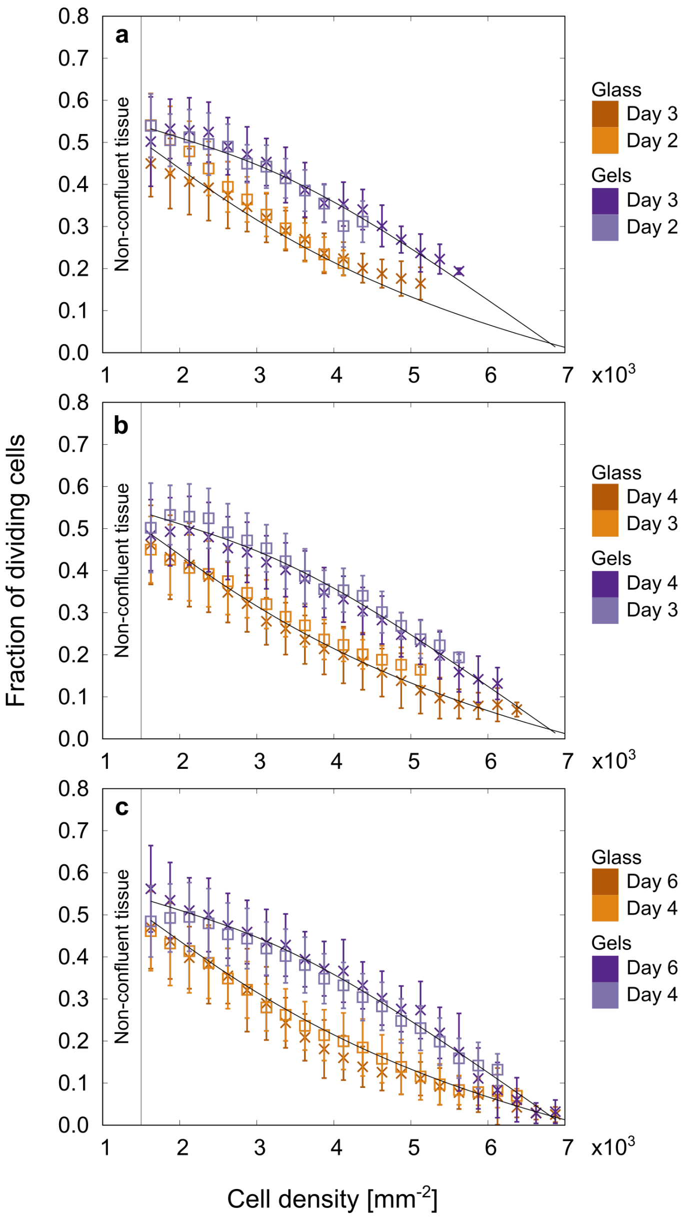

**Fig. S2. Division of MDCK-II cells in confluent tissues and its relation to the cell local environment: Highlight on time evolution.** Highlight on the time evolution of the fraction of dividing cells as a function of the local environment. **a** days 2 and 3 after seeding, **b** days 3 and 4 after seeding, and **c** days 4 and 6 after seeding. The solid line is the fitted cell-level model on all four time points together. Early days (day 2 and day 3) do not span the whole density range, which is only accessible at day 6.

#### S2. Derivation of the cell-level model mean growth time

We start this derivation from the definition of $\mathcal{R}(\rho)$, i.e.

$$\mathcal{R}(\rho) = \left\langle \frac{N_d}{N_g + N_d} \right\rangle, \qquad [1]$$

where $N_g$ and $N_d$ are the number of cell in the *growing* and *dividing* phase respectively in a constant cell density $\rho$. If we assume the cell to spend a time $\tau_g$ per cell cycle in the *growing* phase and a time $\tau_d$ in *dividing* phase and cells to behave sufficiently independently, this leads to

$$\mathcal{R}(\rho) = \frac{\langle \tau_d \rangle}{\langle \tau_g \rangle + \langle \tau_d \rangle}. \qquad [2]$$

We assume that the density-dependence is solely carried by the time spent in the *growing phase* $\tau_g$. Our main task is therefore to provide an analytical model to obtain a mean density-dependent value for the time spent growing $\langle \tau_g \rangle$, whereas $\langle \tau_d \rangle$ will be assumed a constant parameter of the overall model.

To model the time $\langle \tau_g \rangle$, we will observe a single cell of size $\sigma(t)$ in a constant local density $\rho$, which will represent the size parameter of that cell. We model $\tau_g$ via a probability distribution $p_g(t)$, which denotes the probability for a cell which has left the division phase and entered into its growing state at time $t = 0$ to not have entered into its next division phase at time $t > 0$. These assumptions allow us to derive an analytical prediction for the mean value of $\langle \tau_g \rangle$ via an intermediate result for the evolution of $p_g(t)$.

**A. Derivation.** For the time evolution of $p_g(t)$, we assume a traditional rate-decay model for the evolution of the probability $p_g(t)$:

$$\frac{dp_g}{dt} = -r_d(\sigma) p_g \qquad [3]$$

where the change in probability to not have divided is proportional to the remaining probability density $p_d(t)$ via a proliferation rate coefficient $r_d(\sigma)$, which we assume to be solely dependent on the current size of the observed cell.

Via a separation of variables in eq. (3), we can solve the differential equation as

$$\begin{aligned} \frac{1}{p_g}\frac{dp_g}{dt} &= -r(\sigma(t)), \\ \frac{d\,\ln(p_g)}{dt} &= -r(\sigma(t)), \\ p_g(t) &= \exp\left(-\int_0^t r_d(\sigma(t'))\,dt'\right). \end{aligned} \qquad [4]$$

Then we can calculate the mean growth time $\langle \tau_g \rangle$ as

$$\langle \tau_g \rangle = \int_0^\infty p(t)\,dt. \qquad [5]$$

Now, to obtain the precise evolution of $p_g(t)$ and of $\langle \tau_g \rangle$, we need to make assumption regarding the evolution of $r_d(\sigma)$ and $\sigma(t)$. Through the main article, we assume the rate $r_d(\sigma)$ to be linear with respect to the cell size $\sigma$ with a slope of $c_d$ starting at a minimum required size $\sigma_h$ for proliferation to occur and therefore defining the homeostatic state. This leads to the following formula

$$r_d(\sigma) = \begin{cases} 0 & \text{if } \sigma \le \sigma_h, \\ c_d(\sigma - \sigma_h). & \text{else} \end{cases} \qquad [6]$$

Furthermore, we assume the time-evolution of the cell size $\sigma(t)$ to be linear starting from an initial mean size $\sigma_0$ to a final size $\sigma_\rho$ with a slope of $c_g$, i.e. $\sigma_\rho = \sigma_0 + c_g t_\rho$. The size saturates at the final value $\sigma_\rho$ controlled by the isotropic pressure within the tissue from cell density. The final size $\sigma_\rho$, and through a self-consistency argument also $\sigma_0$, are then controlled by the local density $\rho$, whereas we assume the growth speed $c_g$ to be a cell-type and system constant and thus a parameter of the model. Overall, these assumptions lead to

$$\sigma(t) = \begin{cases} \sigma_0 + c_d t & \text{if } t < t_\rho, \\ \sigma_\rho & \text{if } t \ge t_\rho. \end{cases} \qquad [7]$$

Plugging all these information together, we get three different regimes in the growth of the cell

- For $\sigma(t) \le \sigma_h$, linear growth with no proliferation.
- For $\sigma_\rho > \sigma(t) > \sigma_h$, the cell grows to be large enough to proliferate, while statistically being able to divide.
- For $\sigma(t) = \sigma_\rho$, the cell size saturates and so does the rate of proliferation, leading to an exponential decay of the remaining probability $p_g(t)$ over time as the cell stays at constant size while awaiting the onset of division.

It must be stated that the second and third points are to be considered only if $\sigma_\rho > \sigma_c$ leading to a stop of division as soon as local density exceeds the target density $\rho_h$. These three different regimes contribute a total of three different terms to the overall formula for the mean value of $\langle \tau_g \rangle$.

For simplified mathematical notation, let $t_h$ denote the time at which $\sigma(t) = \sigma_h$, and $t_\rho$ be the time, at which the condition $\sigma(t) = \sigma_\rho$ first holds. Then, for simpler notation, let us denote the integral in eq. (4) by $I(t)$ which reads

$$I(t) = \int_0^t r_d(\sigma(s))\, ds = \begin{cases} 0 & \text{if } t \leq t_h, \\ \frac{c_d}{2c_g}(\sigma(l) - \sigma_h)^2 & \text{if } t_h < t \leq t_\rho, \\ \frac{c_d}{2c_g}(\sigma_\rho - \sigma_h)^2 + c_d(t - t_\rho)(\sigma_\rho - \sigma_h) & \text{if } t_\rho < t. \end{cases}$$

We then arrive at the relation

$$p_g(t) = \exp\left(-I(t)\right), \qquad [8]$$

and consequently

$$\langle \tau_g \rangle = \int_0^\infty \exp\left(-I(t)\right)\, dt. \qquad [9]$$

For $\sigma_\rho < \sigma_h$, the integral on the right hand side diverges and $\langle \tau_g \rangle \to \infty$ as $I(t) = 0, \forall t \geq 0$. This agrees with our understanding that cells cannot proliferate if they are not able to attain a certain minimum size.

Let us instead assume that $\sigma_\rho \geq \sigma_h$, then we can solve above integral, using the parameters $t_r = 1/\sqrt{c_d c_g}$ and $\sigma_r = \sqrt{c_g/c_d}$, and $\Delta\sigma = \sigma_\rho - \sigma_h$ to simplify notation:

$$\langle \tau_g \rangle = \int_0^\infty \exp\left(-I(t)\right)\, dt \qquad [10]$$

$$= t_1 + \int_{t_1}^{t_\rho} \exp\left(-\frac{(\sigma(s) - \sigma_h)^2}{2\sigma_r^2}\right) ds + \int_{t_\rho}^\infty \exp\left(-\frac{(\sigma_f - \sigma_h)^2}{2\sigma_r^2} - \frac{(t - t_\rho)}{t_r}\frac{(\sigma_f - \sigma_h)}{\sigma_r}\right) ds \qquad [11]$$

$$= t_r \times \frac{\sigma_h - \sigma_0}{\sigma_r} + \int_{t_1}^{t_\rho} \exp\left(-\frac{(\sigma(s) - \sigma_h)^2}{2\sigma_r^2}\right) ds + \exp\left(-\frac{(\sigma_f - \sigma_h)^2}{2\sigma_r^2}\right) \int_{t_\rho}^\infty \exp\left(-r_d(\sigma_\rho)(t - t_\rho)\right) ds \qquad [12]$$

$$= t_r \times \frac{\sigma_h - \sigma_0}{\sigma_r} + \frac{t_r}{\sigma_r} \int_{\sigma_h}^{\sigma_\rho} \exp\left(-\frac{(o - \sigma_h)^2}{2\sigma_r^2}\right) do + \exp\left(-\frac{(\Delta\sigma)^2}{2\sigma_r^2}\right) \left[-\frac{1}{r_d(\sigma_\rho)} \exp\left(-r_d(\sigma_\rho)(t - t_\rho)\right)\right]_{t=t_\rho}^{t=\infty} \qquad [13]$$

$$= t_r \times \frac{\sigma_h - \sigma_0}{\sigma_r} + t_r\sqrt{2\pi} \int_{\sigma_h}^{\sigma_\rho} \mathcal{N}(\sigma_h, \sigma_r)[o]\, do + \exp\left(-\frac{(\Delta\sigma)^2}{2\sigma_r^2}\right) \frac{1}{r_d(\sigma_\rho)} \qquad [14]$$

$$= t_r \times \frac{\sigma_h - \sigma_0}{\sigma_r} + t_r \times \sqrt{\pi/2} \times \text{erf}\left(\frac{\Delta\sigma}{\sqrt{2}\sigma_r}\right) + t_r \times \frac{\sigma_r}{\Delta\sigma} \exp\left(-\frac{(\Delta\sigma)^2}{2\sigma_r^2}\right) \qquad [15]$$

$$= t_r \left[\frac{\sigma_h - \sigma_0}{\sigma_r} + \sqrt{\pi/2} \times \text{erf}\left(\frac{\Delta\sigma}{\sqrt{2}\sigma_r}\right) + \frac{\sigma_r}{\Delta\sigma} \exp\left(-\frac{(\Delta\sigma)^2}{2\sigma_r^2}\right)\right] \qquad [16]$$

where $\mathcal{N}(m, \nu)$ denotes the normal distribution of mean $m$ and standard deviation $\nu$, and erf denotes the associated error function. The formula in eq. (16) is the analytical prediction for the mean growth time, which depends on the parameters $t_r$, $\sigma_r$, $\sigma_h$, and $\sigma_0$ (where $t_r$ and $\sigma_r$ are a reparametrization of $c_d$ and $c_g$). However, all these parameters are not independent. Indeed, $\sigma_0$ can be related to the other parameters. One can assume $\sigma_0 = \langle \sigma \rangle / 2$ with $\langle \sigma \rangle$ being the average cell size at time of proliferation.

Following this connection, we can calculate a self-consistency condition for the mean size at time of division $\langle \sigma \rangle$ to determine $\sigma_0$ for the case $\sigma_0 \leq \sigma_h < \sigma_\rho$:

$$\langle \sigma \rangle = \int_0^\infty \left(-\frac{dp_g}{dt}\right) \sigma(s)\, ds \qquad [17]$$

$$= \int_{t_h}^{t_\rho} -\frac{dp_g}{dt} \sigma(s)\, ds + p_g(t_\rho)\sigma_\rho \qquad [18]$$

$$= -\left[\sigma(s) p_g(s)\right]_{s=t_h}^{s=t_\rho} + \int_{t_1}^{t_\rho} p(s) \frac{d\sigma}{dt}\, ds + p_g(t_\rho)\sigma_\rho \qquad [19]$$

$$= \sigma_h p_g(t_h) - \sigma_\rho p_g(t_\rho) + \sigma_r \sqrt{\frac{\pi}{2}} \text{erf}\left(\frac{(\sigma_\rho - \sigma_h)}{\sqrt{2}\sigma_r}\right) + p_g(t_\rho)\sigma_\rho \qquad [20]$$

$$= \sigma_h + \sigma_r \sqrt{\frac{\pi}{2}} \text{erf}\left(\frac{\Delta\sigma}{\sqrt{2}\sigma_r}\right) \qquad [21]$$

where $p_g(t_1) = 1$ has been used. This final result can be used to obtain $\sigma_0 = \langle\sigma\rangle/2$ and plug the final expression into eq. (16). One obtains

$$\langle\tau_g\rangle = t_r \left[\frac{\sigma_h}{2\sigma_r} + \sqrt{\pi/8} \times \mathrm{erf}\left(\frac{\Delta\sigma}{\sqrt{2}\sigma_r}\right) + \frac{\sigma_r}{\Delta\sigma}\exp\left(-\frac{(\Delta\sigma)^2}{2\sigma_r^2}\right)\right] \tag{22}$$

Equation (22) is now fully parameterized by $\sigma_h$, $t_r$, and $\sigma_r$ (and through those $c_d$ and $c_g$). The density dependence is encoded in the parameter $\Delta\sigma = \sigma_\rho - \sigma_h$ through the density dependence of the target size $\sigma_\rho$.

Due to the need for an upper limit that is up or above $1/\rho$ wherever the tissue is confluent, we opted for the following function as an upper limit for the cell size as a function of the surrounding density $\rho$:

$$\sigma_\rho = \frac{2 - (\rho\sigma_h)^{0.1}}{\rho} \tag{23}$$

The term $2 - (\rho\sigma_h)^{0.1}$ was chosen, because it is only slightly above 1 for a very long time, whereas at low densities it increases to about 2. This is necessary, because $\sigma_{\rho_h} = \sigma_h$ needs to hold, but $\sigma_h > 1/\rho$ is required everywhere below $\rho_h$ for the model to remain self-consistent. As shown in fig. A1, the chosen upper boundary $\sigma_\rho$ according to eq. (23) keeps the predicted mean size at time of divison, the maximum size of the cell and the minimum required size for confluence self-consistent while approximately predicting the correct density at which confluence of colonies is lost. We therefore believe the expression in eq. (23) to be a good approximation of the real behavior observed in tissues as confirmed by the accuracy of our fits.

Adding to $\mathcal{R}$ the parameter $\langle\tau_d\rangle$ — the constant duration the division phase — one can give a full prediction for the behavior of $R$.

**B. Low-density considerations.** At $\rho \to 0$, we may for other choices of $\sigma_\rho$ observe such a fast growth, that $\sigma_0 > \sigma_h$, let us consider the impact of this on the self-consistency argument for $\sigma_0$. First:

$$I(t) = \int_0^t r_d(\sigma(s))\,ds = \begin{cases} \frac{(\sigma(l)-\sigma_h)^2-(\sigma_0-\sigma_h)^2}{2\sigma_r^2} & \text{if } t \le t_\rho, \\ \frac{(\sigma_\rho-\sigma_h)^2-(\sigma_0-\sigma_h)^2}{2\sigma_r^2} + \frac{1}{t_r\sigma_r}(t-t_\rho)(\sigma_\rho-\sigma_h) & \text{if } t_\rho < t. \end{cases}$$

Plugging this into the derivation of the mean cell size at time of division, we get:

$$\langle\sigma\rangle = \int_0^\infty \left(-\frac{dp_g}{dt}\right)\sigma(s)\,ds \tag{24}$$

$$= \int_0^{t_\rho} -\frac{dp_g}{dt}\sigma(s)\,ds + p_g(t_\rho)\sigma_\rho \tag{25}$$

$$= -\left[\sigma(s)p_g(s)\right]_{s=0}^{s=t_\rho} + \int_0^{t_\rho} p(s)\frac{d\sigma}{dt}\,ds + p_g(t_\rho)\sigma_\rho \tag{26}$$

$$= \sigma_0 p_g(0) - \sigma_\rho p_g(t_\rho) + \sigma_r\sqrt{\frac{\pi}{2}}\exp\left(-\frac{(\sigma_0-\sigma_h)^2}{2\sigma_r^2}\right)\left[\mathrm{erf}\left(\frac{(\sigma_\rho-\sigma_h)}{\sqrt{2}\sigma_r}\right) - \mathrm{erf}\left(\frac{(\sigma_0-\sigma_h)}{\sqrt{2\sigma_r^2}}\right)\right] + p_g(t_\rho)\sigma_\rho \tag{27}$$

$$= \sigma_0 + \sigma_r\sqrt{\frac{\pi}{2}}\exp\left(-\frac{(\sigma_0-\sigma_h)^2}{2\sigma_r^2}\right)\left[\mathrm{erf}\left(\frac{\Delta\sigma}{\sqrt{2}\sigma_r}\right) - \mathrm{erf}\left(\frac{\sigma_0-\sigma_h}{\sqrt{2}\sigma_r}\right)\right]. \tag{28}$$

Then the self-consistency argument amounts to

$$\sigma_0 = \sigma_r\sqrt{\frac{\pi}{2}}\exp\left(-\frac{(\sigma_0-\sigma_h)^2}{2\sigma_r^2}\right)\left[\mathrm{erf}\left(\frac{\Delta\sigma}{\sqrt{2}\sigma_r}\right) - \mathrm{erf}\left(\frac{\sigma_0-\sigma_h}{\sqrt{2}\sigma_r}\right)\right]. \tag{29}$$

This condition for fixed $\sigma_r$ and $\sigma_h$ has at most one solution $\sigma_0$, as the left hand side is strictly monotonously increasing and the right hand side strictly monotonously decreasing in $\sigma_0$. If a solution exists, where both sides are equal, this results in a density-independent limit for $\sigma_0$ at extremely low densities, denoted by:

$$\sigma_h < \sigma_0 \le \sigma_r\sqrt{\frac{\pi}{2}}$$

which can only ever be satisfied, if $\sigma_h < \sigma_r\sqrt{\frac{\pi}{2}}$, but for this scenario, there will be a density cutoff, where the mean cell size at time of division does not increase any further. Therefore, only if $\sigma_r > \sqrt{\frac{2}{\pi}}\sigma_h$ can this issue even arise and even then, it will only be an issue for very low densities close to the single-cell scenario.

As our data in the low-density limit is limited and our data does not satisfy the necessary conditions in the relevant density interval, we have decided to only consider the case $\sigma_0 \le \sigma_h < \sigma_\rho$ for our derivation employed in the main manuscript.

Still we provide a plot of the characteristic dependence of the curves $\tau_g$ and $\langle\sigma\rangle$ on the parameters $t_r$ and $\sigma_r$ in the high- and low-density limits where applicable in fig. S3. The effect on $\mathcal{R}$ is presented in the main manuscript.

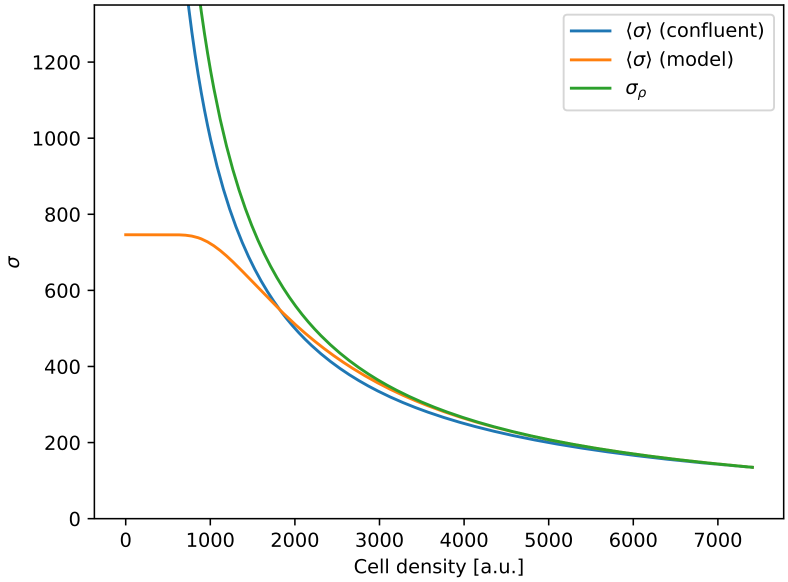

**Fig. A1. Size constraints and confluency in the Cell-level model.** The plot of the upper boundary $\sigma_\rho$ of cell size according to the cell-level model, the mean expected cell size at time of division ($\langle\sigma\rangle_{\mathrm{model}}$)according to the model predictions and the necessary mean size $\langle\sigma\rangle_{\mathrm{confluence}} = 1/\rho$, that is required for confluence to be maintained. The plot is obtained for a glass-substrate model system as according to the fit parameters in table S1. The upper boundary is only slightly above the mean required size, which is the desired behavior of such a boundary, supporting our choice of $\sigma_\rho$. Notably, we also arrive at a crossover at aroung $\rho \approx 1800\,\mathrm{cells}/\mathrm{mm}^2$, where confluence would be lost, which is very close to the observed loss of confluence at about$1500\,\mathrm{cells}/\mathrm{mm}^2$ in experiments. This further supports the appropriate nature of the model choices made.

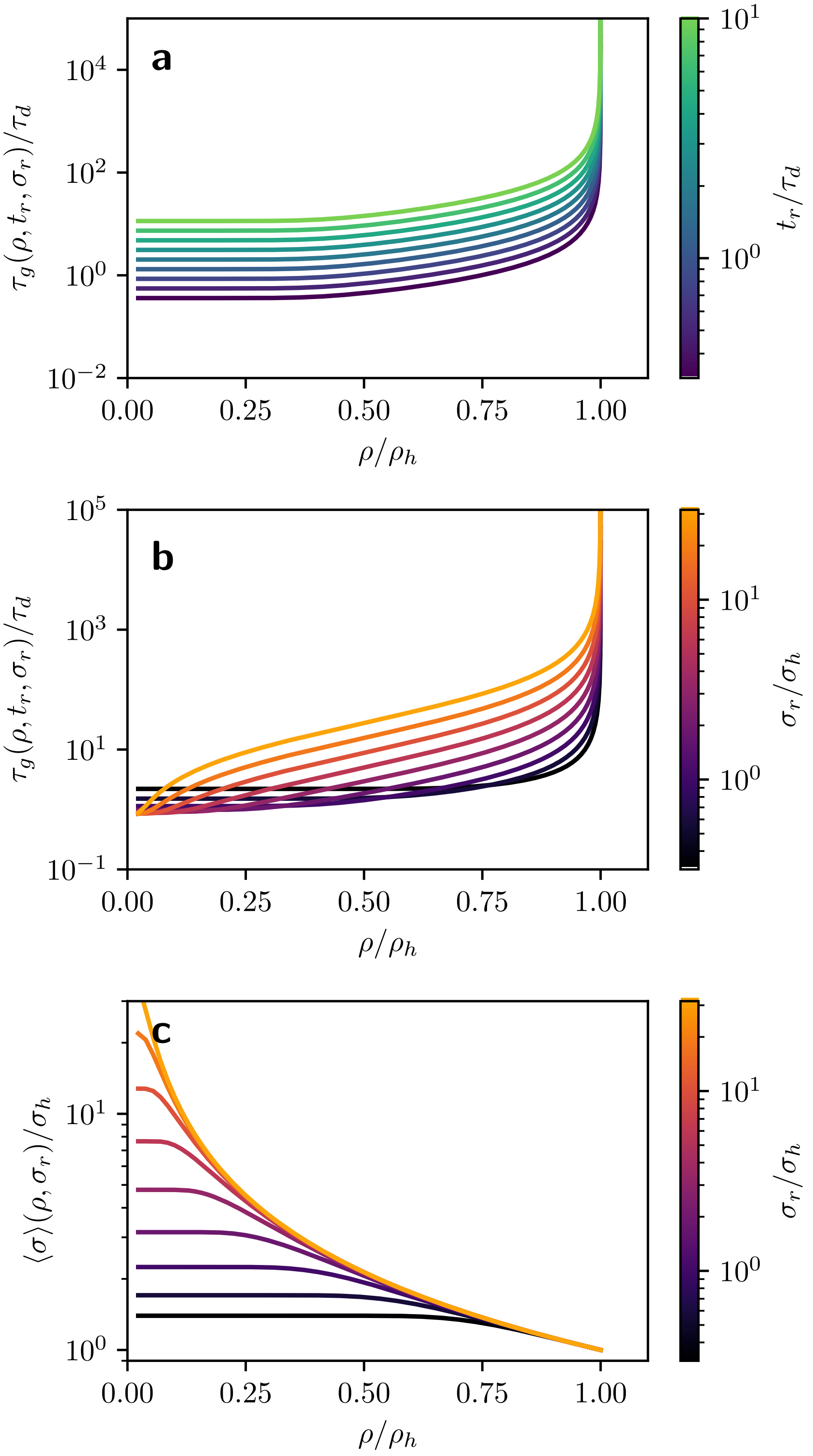

**Fig. S3. Influence of the parameters $t_r$ and $\sigma_r$ on the mean growth time and mean area at time of division cells.** Evolution of the mean growth time $\tau_g$ as a function of the density (**a**,**b**) with the effect of $t_r$ (**a**) and $\sigma_r$ (**b**) being color-coded. The evolution of the mean cell size $\langle\sigma\rangle$ at time of division as a function of the density is plotted in **c** with the effect of $\sigma_r$ on the evolution being color-coded. As stated in the main article, $\sigma_r$ mainly carries the changes of curvature through the inflection point/saturation cutoff of $\tau_g$.

**C. Time- and Area statistics according to the cell-level model.** The knowledge of eq. (8) gives access to the distribution of growth time and dividing cell area. Indeed, one can show that the probability to not have divided at time $t$ reads

$$p_g(t) = \begin{cases} 1, & \text{if } t < t_h = \frac{\sigma_h - \sigma_0}{c_g} \\ \exp\left(-\frac{c_d}{2c_g}(\sigma(t) - \sigma_h)^2\right), & \text{if } t_h < t < t_\rho = \frac{\sigma_\rho - \sigma_0}{c_g} \\ \exp\left(-\frac{c_d}{2c_g}(\sigma_\rho - \sigma_h)^2 - r_0(t - t_\rho)(\sigma_\rho - \sigma_h)\right), & \text{if } t_\rho < t \end{cases} \quad [30]$$

which provides a plateau for $\sigma(t) < \sigma_h$, a Gaussian decrease for $\sigma_h < \sigma(t) < \sigma_\rho$, and an exponential decrease beyond, therefore agreeing with the results from Smith and Martin (4).

We also want to investigate the distribution of the cell size $\sigma(t)$ in the division phase. For this purpose, let $T(\sigma)$ denote the time at which the cell attains size $\sigma$ during its growth, where $\sigma \leq \sigma_\rho$ or else this moment does not exist. Then the probability for the cell to enter the division state with size $\sigma = \sigma_\rho$ is

$$p_d(\sigma = \sigma_\rho) = p_g(T(\sigma_\rho)) = p_g(t_\rho).$$

Since $\sigma_h$ is the minimum size and $\sigma_\rho$ the maximum size of a dividing cell, we can further deduce:

$$p_d(\sigma > \sigma_\rho) = p_d(\sigma < \sigma_h) = 0$$

For $\sigma_h < \tilde{\sigma} < \sigma_\rho$ the relation between size and time $\sigma(t) = c_g(t - t_1) + \sigma_h$ leads us to

$$p_d(\sigma < \tilde{\sigma}) = 1 - p_g(T(\tilde{\sigma})) = 1 - p_g\left(\frac{\tilde{\sigma} - \sigma_h}{c_g} + t_h\right). \quad [31]$$

Let $\alpha_d$ denote the distribution of cell area (i.e. probability distribution) of cells entering the division part of the proliferation cycle. Then we arrive at

$$\begin{aligned} \alpha_d(\tilde{\sigma}) &= \partial_{\tilde{\sigma}} p_d(\sigma < \tilde{\sigma}) & [32] \\ &= -\frac{1}{c_g}\partial_t p(T(\tilde{\sigma})) & [33] \\ &= \frac{r_d(\tilde{\sigma})}{c_g} p(T(\tilde{\sigma})) & [34] \\ &= \frac{r_d(\tilde{\sigma})}{c_g} \exp\left(-\frac{c_d}{2c_g}(\tilde{\sigma} - \sigma_h)^2\right) & [35] \\ &= \frac{1}{\sigma_r}\frac{\tilde{\sigma} - \sigma_h}{\sigma_r} \exp\left(-\frac{(\tilde{\sigma} - \sigma_h)^2}{2\sigma_r^2}\right) & [36] \end{aligned}$$

As a result, the entire density distribution in division can be written as

$$\alpha_d(\sigma) = p(t_\rho)\delta(\sigma - \sigma_\rho) + \begin{cases} 0 & \text{if } \sigma < \sigma_h \\ \frac{1}{\sigma_r}\frac{\sigma - \sigma_h}{\sigma_r} \exp\left(-\frac{(\sigma - \sigma_h)^2}{2\sigma_r^2}\right) & \text{if } \sigma_h \leq \sigma < \sigma_\rho \\ 0 & \text{if } \sigma \geq \sigma_\rho \end{cases} \quad [37]$$

#### S3. Effect of $t_r$ and $\sigma_r$ on growth duration and division statistics of isolated cells

Let us take a look at the limit $\rho \to 0$ for the scenario $\sigma_0 \leq \sigma_h$ to investigate the effect the characteristic effects of $t_r$ and $\sigma_r$. First, we note that:

$$\sigma_\rho = \frac{2 - (\rho\sigma_h)^{0.1}}{\rho} \to \infty \qquad [38]$$

and consequently

$$\Delta\sigma \to \infty \qquad [39]$$

From our derivation in eq. (21), we can conclude, that the mean size of cells at time of division in the low density limit should then follow:

$$\langle\sigma\rangle \approx \sigma_h + \sigma_r\sqrt{\frac{\pi}{2}} \qquad [40]$$

meaning that at low densities, $\sigma_r$ will control the excess average size of a cell above $\sigma_h$ at time of division in a linear fashion. This excess-scaling-link even holds true considering the low-density limit in eq. (29) despite the obvious linearity being lost. I.e. even in the low-density derivation, the mean size of a single cell at time of division increases with $\sigma_r$ and for small values of $\sigma_r$ above the necessary threshold, this relation is approximately linear.

Let us now instead consider the mean growth time $\langle\tau_g\rangle$ in this limit. From eq. (22) we then derive:

$$\begin{aligned}\langle\tau_g\rangle(\rho \to 0) =& t_r\left[\frac{\sigma_h}{2\sigma_r} + \sqrt{\pi/8} \times \operatorname{erf}\left(\frac{\Delta\sigma}{\sqrt{2}\sigma_r}\right) + \frac{\sigma_r}{\Delta\sigma}\exp\left(-\frac{(\Delta\sigma)^2}{2\sigma_r^2}\right)\right] & [41]\\ \approx& t_r\left[\frac{\sigma_h}{2\sigma_r} + \sqrt{\pi/8}\right] & [42]\end{aligned}$$

This means, that $t_r$ and $\sigma_r$ control the mean time between divisions at low density together with the critical size threshold $\sigma_h$. As we can determine $\sigma_r$ from the distribution of cell sizes at the beginning of the division phase, we can then employ that knowledge to derive the time constant $t_r$ from the distribution of growth times.

Looking at the probability of cells to not have divided by time $t$ (eq. (30)), we see:

$$p_g(t) = \begin{cases} 1, & \text{if } t < t_h = t_r \times \frac{\sigma_h - \sigma_0}{\sigma_r} \\ \exp\left(-\frac{(\sigma(t)-\sigma_h)^2}{2\sigma_r^2}\right), & \text{if } t_1 < t < t_\rho = t_r \times \frac{\sigma_\rho - \sigma_0}{\sigma_r} \\ \exp\left(-\frac{(\sigma_\rho-\sigma_h)^2}{2\sigma_r^2} - c_d(t - t_\rho)(\sigma_\rho - \sigma_h)\right), & \text{if } t_\rho < t \end{cases} \qquad [43]$$

$$= \begin{cases} 1, & \text{if } t < t_h \\ \exp\left(-\frac{(t-t_h)^2}{2t_r^2}\right), & \text{if } t_h < t < t_\rho \\ \exp\left(-\frac{(\sigma_\rho-\sigma_h)^2}{2\sigma_r^2}\right) \times \exp\left(-\frac{\sigma_\rho-\sigma_h}{\sigma_r} \times \frac{t-t_\rho}{t_r}\right), & \text{if } t_\rho < t \end{cases}. \qquad [44]$$

From this, we conclude, that $t_r$ is a scale factor, of the time statistics during the entire growth phase throughout the colony, linearly scaling up and down the distribution of all growth times, also affecting the slope of the distribution in its Gaussian-regime and the characteristic time scale in the exponential decay after saturation.

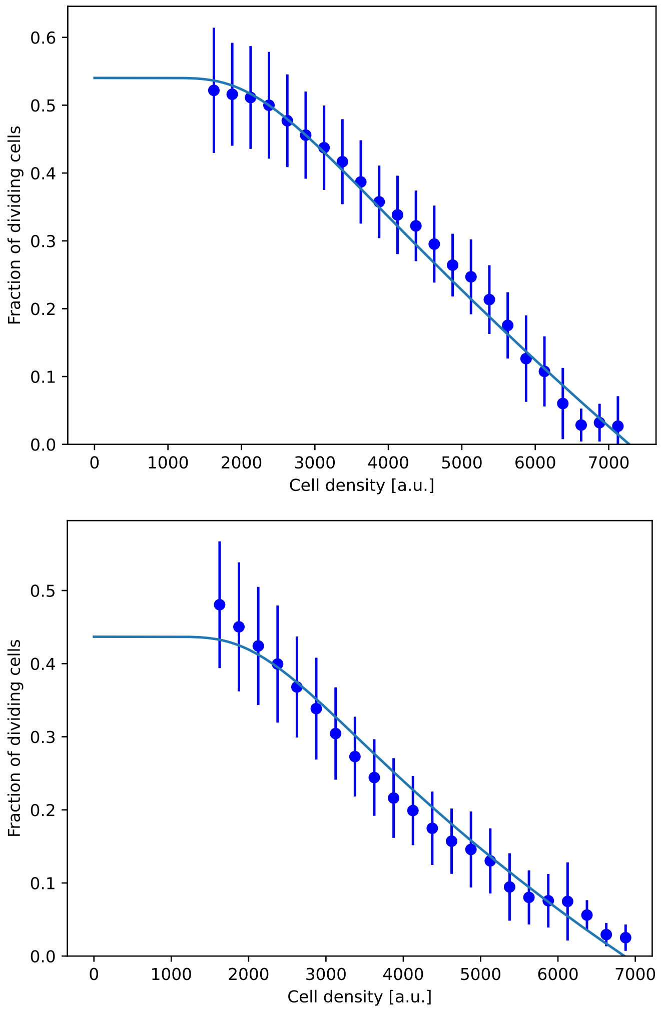

**Fig. S4. Cell-level model fits to experimental data with constrained reference homeostatic density.** Plot of the fits performed for table S1 with $\rho_{h,gel} = 7280\,\mathrm{cells/mm^2}$ (top) and $\rho_{h,glass} = 6860\,\mathrm{cells/mm^2}$ (bottom). The fit curves without fixed reference homeostatic densities $\rho_h$ are presented in the main manuscript.

**Table S1. Fitting parameters to the MDCK-II experiments presented in the relevant equation in the main script. The first two rows also used $\rho_h$ as a fit parameter, whereas in the last two rows, the value of $\rho_h$ was fixed to the respective homeostatic densities reported in a previous publication (1). For our calculations of characteristic statistics we will rely on the first two rows.**

| **Substrate** | $t_r/\tau_d$ [-] | $\sigma_r$ [mm$^2$] | $\rho_h = \sigma_h^{-1}$ [cells/mm$^2$] |
|---|---|---|---|
| Free fit of $\rho_h$, $t_r/\tau_d$ and $\sigma_r$ | | | |
| $11\,\mathrm{kPa}$ **gels** | $0.90$ | $1.8170 \times 10^{-4}$ | $6837(62)$ |
| **glass** | $0.91$ | $5.1471 \times 10^{-4}$ | $7407(56)$ |
| Fixed $\rho_h$ to reference value (1) | | | |
| $11\,\mathrm{kPa}$ **gels** | $0.87$ | $2.5450 \times 10^{-4}$ | 7280 |
| **glass** | $1.30$ | $2.5176 \times 10^{-4}$ | 6860 |

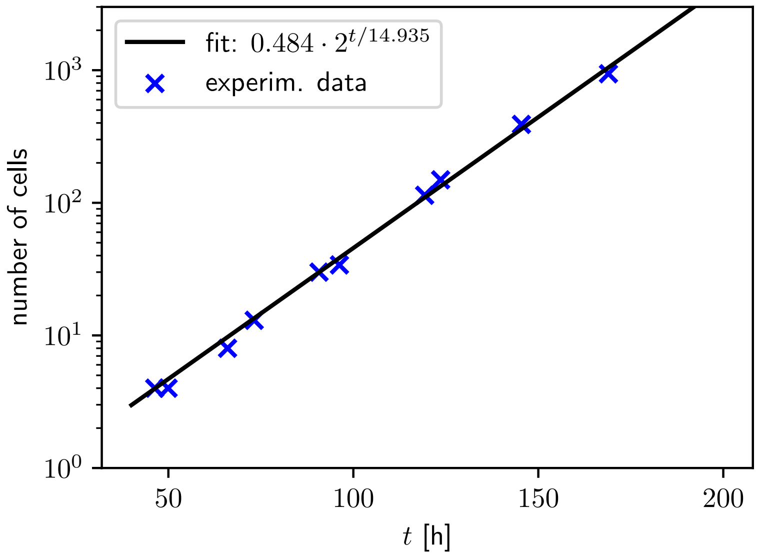

**Fig. S5. Evolution of the number of cells in a low-density configuration on glass substrate.** Experimental results of the number of observed cells in a small colony (i.e. almost isolated cells over time (blue) as well as a fit with an exponential curve (black), yielding a full doubling/cell cycle duration of $\approx 15\,\mathrm{h}$. (5)

#### S4. Validating the cell-level model's predictions against experimental data

According to a publication by Devany *et al.*, the total division cycle time on glass is

$$\tau_{s+G_2+M} = 10\,\mathrm{h}.$$

Based on our fit of the doubling time of low cell counts on glass (see fig. S5):

$$\tau_g + \tau_d = 15\,\mathrm{h}$$

According to the derivation presented in eq. (29):

$$\langle\sigma\rangle \approx \sigma_h + \sigma_r\sqrt{\frac{\pi}{2}} \quad [45]$$

$$\approx \rho_h^{-1} + \sigma_r\sqrt{\frac{\pi}{2}} \quad [46]$$

and according to eq. (42):

$$\langle\tau_g\rangle(\rho \to 0) \approx t_r\left[\frac{\sigma_h}{2\sigma_r} + \sqrt{\pi/8}\right] \quad [47]$$

For the mean area of an isolated cell at time of division using the results from table S1, for glass this leaves us with

$$\langle\sigma\rangle_{\mathrm{glass}} = 7407^{-1} + 5.1471 \times 10^{-4}\sqrt{\frac{\pi}{2}}\,\mathrm{\mu m}^2 \quad [48]$$

$$= 780.1\,\mathrm{\mu m}^2 \quad [49]$$

and for 11 kPa gels:

$$\langle\sigma\rangle_{\mathrm{gel}} = 6837^{-1} + 1.8170 \times 10^{-4}\sqrt{\frac{\pi}{2}}\,\mathrm{\mu m}^2 \quad [50]$$

$$= 374.0\,\mathrm{\mu m}^2 \quad [51]$$

i.e. an isolated cell on glass is expected to be about 109 % larger than a similar cell grown on a gel substrate when it starts to divide. This is not as drastic a difference as observed in previous results, where the isolated glass cells were larger by a factor of almost 6 on soft substrates (6), but still in line with the observation of isolated cells on stiff gels being approximately 30 % smaller (7).

Plugging in the results for glass into the lifetime expectations, this leaves us with:

$$\tau_g = t_r\left[\frac{1}{2\sigma_r\rho_h} + \sqrt{\pi/8}\right] \quad [52]$$

$$= t_r\left[\frac{1}{2 \cdot 5.1471 \times 10^{-4} \cdot 7407} + \sqrt{\pi/8}\right] \quad [53]$$

$$= 0.757806 \cdot t_r \Leftrightarrow t_r = 1.32\tau_g \quad [54]$$

We can estimate, that the duration of the s-phase will be about 2/3 of the full division time of 10 h. Due to the finite time of EdU integration as well as the thresholding applied to the EdU signal in post-processing, the stained population should account for 2 more hours than that, leaving us with an estimate of

$$\tau_d \approx 8.67\,\mathrm{h}.$$

Assuming the remaining time to be $\tau_g$, then leaves us with

$$\tau_g \approx 6.33\,\mathrm{h}$$

and consequently

$$t_r \approx 8.29\,\mathrm{h}$$

which leaves us with

$$\frac{t_r}{\tau_d} = 0.829$$

which agrees nicely with our result of 0.91, when fitting the $R$ curve with our cell-level model.

Overall, the independently obtained division phase durations and doubling times of cells on glass appear to agree nicely with our fit results for the ratio of $\frac{t_r}{\tau_d}$ corroborating our results.

#### S5. Issues with simpler models

In the context of cell proliferation, other models than the one introduced in the main article has been proposed (8–12). We will now perform the calculations for the growing time for those other models.

**A. Deterministic proliferation.** An option discussed in the literature is the case of instantaneous proliferation above a given size threshold. This is equivalent to $c_d \to \infty$ for $\sigma > \sigma_h$. We will use this scheme in the mathematical framework introduced earlier in this supplementary informations to extract the resulting $\langle \tau_g \rangle$ and $\mathcal{R}$.

The assumption $c_d \to \infty$ for $\sigma > \sigma_h$ can also be written as

$$p_g(t) = \begin{cases} 1 & t < t_h, \\ 0 & t \geq t_h. \end{cases} \quad [55]$$

Thus, the cell always divides at $\sigma = \sigma_h$ and we can simplify the expression of $\sigma_0$ to $\sigma_0 = \sigma_h/2$. Therefore, the intermediate integral $I(t) = 0$ for $t < t_h$ while diverging to infinity beyond the time $t_h$. One therefore obtains the following average time spent in the growth phase

$$\langle \tau_g \rangle = \frac{\sigma_h - \sigma_0}{c_g}, \quad [56]$$

$$= \frac{\sigma_h}{2c_g}. \quad [57]$$

[58]

This result only holds for $\rho \leq \rho_h$ and otherwise diverges. As a consequence, the ratio of proliferating cells writes

$$\mathcal{R} = \frac{\langle \tau_d \rangle}{\langle \tau_d \rangle + \frac{\sigma_h}{2c_g}}, \quad [59]$$

for $\rho \leq \rho_h$ and $\mathcal{R} = 0$ for $\rho > \rho_h$. The value is therefore density-independent, and does not possess any further degrees of freedom that are necessary to capture mechanosensitive properties (see fig. S6 for details of shape and behavior). As a conclusion, this model cannot capture any of the result in the main article.

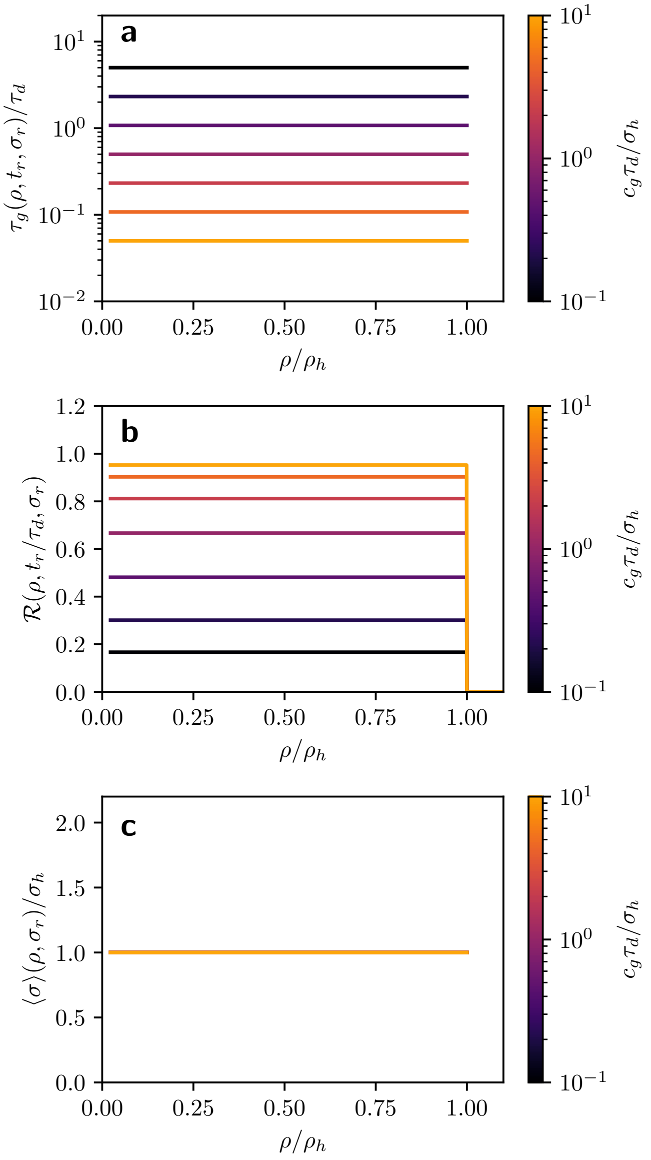

**Fig. S6. Fraction of dividing cells as a function of the cell density in the case of a deterministic (instantaneous) division protocol.** (a) the value of $\tau_g$ as a function of the only parameter $c_g$. (b) The ratio of dividing cells $\mathcal{R}$ as a depending on the parameter $c_g$. Only the magnitude of the fraction of dividing cells is controllable, the shape cannot be changed. (c) The mean size of a cell at time of division. This parameter is independent of the parameter $c_g$.

 

**B. Constant proliferation rate.** Another model is one of constant proliferation rate, instead of the increasing rate presented in the main article. We again base our derivation on a constant growth

$$\sigma(t) = \begin{cases} \sigma_0 + c_g t & t < t_\rho \\ \sigma_\rho & t \geq t_\rho \end{cases} \qquad [60]$$

where we assume a simple size limit relation $\sigma_\rho = 1/\rho$.

Let us now assume a proliferation rate that is initially zero as long as the cell is below $\sigma_h$ in size and then a constant value of $\alpha$, i.e.

$$r_d(\sigma) = \begin{cases} 0 & \sigma < \sigma_h, \\ \alpha & \sigma \geq \sigma_h. \end{cases} \qquad [61]$$

The intermediate integral $I(t)$ therefore becomes

$$I(t) = \int_0^t r_d(\sigma(s))\, ds = \begin{cases} 0 & t < t_h, \\ \alpha(t - t_h) & t \geq t_1 \end{cases}, \qquad [62]$$

and the growth time duration is

$$\begin{aligned} \langle \tau_g \rangle &= \int_0^\infty \exp\left(-I(l)\right)\, dt & [63] \\ &= \frac{\sigma_h - \sigma_0}{c_g} + \int_{t_h}^\infty e^{-\alpha(t - t_h)}\, dt & [64] \\ &= \frac{\sigma_h - \sigma_0}{c_g} + \frac{1}{\alpha} & [65] \end{aligned}$$

Following the procedure developed earlier in this document, we compute the size $\langle \sigma \rangle$ to extract the expression of $\sigma_0$

$$\begin{aligned} \langle \sigma \rangle &= \int_0^\infty -\frac{dp_g}{dt}\sigma(s)\, ds & [66] \\ &= \int_{t_h}^{t_\rho} -\frac{dp_g}{dt}\sigma(s)\, ds + p(t_\rho)\sigma_\rho & [67] \\ &= p(t_h)\sigma_h - p(t_\rho)\sigma_\rho + \int_{t_h}^{t_\rho} p_g(s) c_g\, ds + p(t_\rho)\sigma_\rho & [68] \\ &= \sigma_h + c_g \int_{t_h}^{t_\rho} \exp\left(-\alpha(t - t_h)\right)\, ds & [69] \\ &= \sigma_h + \frac{c_g}{\alpha}\left\{1 - \exp\left[-\alpha(t_\rho - t_h)\right]\right\} & [70] \\ &= \sigma_h + \frac{c_g}{\alpha}\left[1 - \exp\left(-\alpha\frac{\sigma_\rho - \sigma_h}{c_g}\right)\right] & [71] \end{aligned}$$

This result exhibits an exponentially decaying dependence on $\rho$ via $\sigma_\rho$, so the overall time $\langle \tau_g \rangle$ depends on the local density. We plug this result into $\sigma_0 = \langle \sigma \rangle / 2$, which leads to

$$\begin{aligned} \langle \tau_g \rangle &= \frac{\sigma_h c - \sigma_0}{c_g} + \frac{1}{\alpha} & [72] \\ &= \frac{\sigma_h}{2c_g} + \frac{1}{\alpha}\left\{1 - \frac{1}{2}\left[1 - \exp\left(-\alpha\frac{\sigma_\rho - \sigma_h}{c_g}\right)\right]\right\} & [73] \\ &= t_r\left\{\frac{\sigma_h}{2\sigma_r} + \frac{1}{2}\left[1 + \exp\left(-\frac{\sigma_\rho - \sigma_h}{\sigma_r}\right)\right]\right\} & [74] \end{aligned}$$

where we used the parameters $t_r = \frac{1}{\alpha}$ and $\sigma_r = \frac{c_g}{\alpha}$.

As before, we also need to consider low densities separately:

$$\langle\sigma\rangle = \int_0^\infty -\frac{dp_g}{dt}\sigma(s)\,ds \quad [75]$$

$$= \int_0^{t_\rho} -\frac{dp_g}{dt}\sigma(s)\,ds + p(t_\rho)\sigma_\rho \quad [76]$$

$$= p(0)\sigma_0 - p(t_\rho)\sigma_\rho + \int_0^{t_\rho} p_g(s)c_g\,ds + p(t_\rho)\sigma_\rho \quad [77]$$

$$= \sigma_0 + c_g\int_{t_1}^{t_\rho} \exp\left(-\alpha(t)\right)\,ds \quad [78]$$

$$= \sigma_0 + \frac{c_g}{\alpha}\left\{1 - \exp\left[-\alpha(t_\rho)\right]\right\} \quad [79]$$

$$= \sigma_0 + \sigma_r\left[1 - \exp\left(-\frac{\sigma_\rho - \sigma_0}{\sigma_r}\right)\right] \quad [80]$$

with a growth time duration of

$$\langle\tau_g\rangle = \int_0^\infty \exp\left(-I(l)\right)\,dt \quad [81]$$

$$= \int_0^\infty e^{-\alpha t}\,dt \quad [82]$$

$$= t_r \quad [83]$$

which can then be used in the before-detailed self-consistency equation to deduce $\sigma_0$.

This leaves us with a slight density dependence in the second (exponential) term of this formula, that manifests in a finite change in $\tau_g$ but also only a limited change in $\mathcal{R}$, which therefore does not converge to zero as $\tau_g$ does not diverge as density approaches $\rho_h$ (see fig. S7 for details of shape and dependence on parameters $t_r$ and $\sigma_r$). Again, this model cannot capture the experimental results.

 

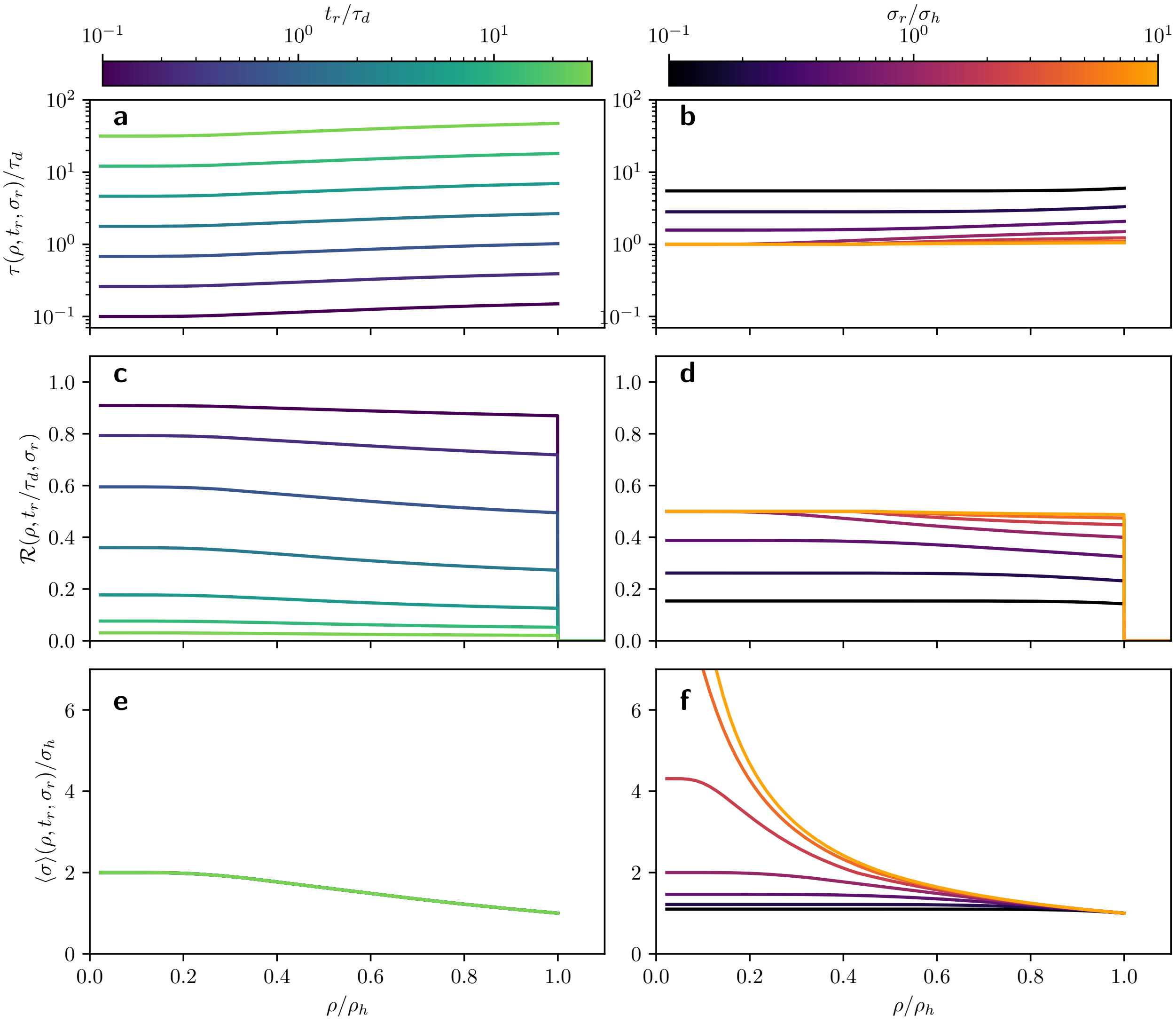

**Fig. S7. Fraction of dividing cells as a function of the cell density in the case of a stochastic division protocol at constant rate.** The behavior of the mean growth time $\tau_g$ (a,b), the fraction of dividing cells $\mathcal{R}$ (c,d) and the mean size of cells at time of division for different values of the parameters $t_r$ (a,c,e) and $\sigma_r$ (b,d,f) respectively (the other parameter is fixed at $\sigma_r = \sigma_h$ and $t_r = \tau_d$ respectively). While $t_r$ sets the scale of $\tau_g$ and $\mathcal{R}$ and the parameter $\sigma_r$ controls the shape of $\tau_g$ and $\mathcal{R}$ in contrast to the instant division model, the density dependence is minor and the curve of $\mathcal{R}$ becomes non-continuous close to the homeostatic density due to the lack of divergence in $\tau_g$. This divergence is essential for reproducing our experimental results and depends on $r(\sigma) \to 0$ as $\sigma \to \sigma_h$, which is the shortcoming of this model.

#### S6. Dissipative Particle Dynamics simulation methods

Dissipative Particle Dynamics (DPD) simulations are an agent-based class of algorithm that have been used successfully in biophysics and which are highly adaptive as it can be seen from the large number of applications in different scientific fields.

In the context of epithelial growth, each cell within the tissue is approximated by two points that are sources and targets of all forces in the system (see fig. 3a in the main manuscript). All forces, viscous or deriving from a potential, are pairwise implying that momentum is conserved in the system. A complete list and mathematical definition of the forces is provided in a previous publication (1). Cells interact with their neighbors via a Lenard-Jones-like potential, allowing for stiff short-range repulsion and soft long-range attraction. Cells are submitted to a viscous drag proportional to the cell velocity measured with respect to the background. A second drag is applied and is proportional to the relative speed between neighboring cells, opposing the relative motion. Within a single cell, during the growth phase, the two building particles repel through a repulsive force mimicking an inner pressure ($\vec{F_g}$), necessary for cell growth and cell division. While in the division phase, the building particles are kept at a constant distance by a stiff harmonic force ($\vec{F_d}$). Also, a random force of uniform angular distribution and normally distributed intensity is applied to all particles and accounts for passive fluctuations in the tissue.

The simulations also account for active ingredients. Cell division is a stochastic process and is made of two phases. The first phase, named the "growth phase", is characterized by a pressure built up within the cells, increasing the distance $l = |\vec{r}_{j,1} - \vec{r}_{j,2}|$ between the two building particles {1,2} of cell $j$. As this distance crosses a threshold $R_{c1}$, the cell as a non-zero probability to enter the second phase or "division phase". The probability is given by

$$\mathcal{P}(l) = \frac{l - R_{c1}}{R_{c2} - R_{c1}}, \qquad [84]$$

where $R_{c2}$ is a second threshold above which the probability is equal to one. Once in the *division* phase, the inner pressure disappears and instead the cell distance is kept at the value it has when entering the *division* phase. While the duration of the growth phase is stochastic and depends on the parameters of the simulations, the duration of the division phase is a parameter and is set to a fixed value.

The final active ingredient of the simulation links cell position and cell shape. The cell shape is obtained by Voronoi Tessellation based on the average distance between the two building points, mimicking the cell nucleus in the simulation. The cell motility is obtained by applying a limited Lloyd optimization step to the cell nuclei position. Specifically, if $d = |\vec{r}_{nuc} - \vec{r}_{cell}|$ where $\vec{r}_{nuc}$ and $\vec{r}_{cell}$ are the nuclei and cell centers respectively, then the nuclei is moved by a distance $\lambda d$ towards the cell center where $\lambda$ is the amplitude of the Lloyd optimization step.

#### S7. Mapping the DPD and cell-level models

One can compute the rate $r(l)$ for the DPD simulations. While this result does not correspond to $r_d(\sigma)$, it will allow us to understand the role of the parameters $R_{c1}$ and $R_{c2}$ in the proliferation rules.

First of all, we assume the simulations to occur in time steps of fixed size $\Delta t$. In each time step, the division probability $P(\ell)$ is calculated, i.e. that a cell will divide according to the following formula

$$P(\ell) = \begin{cases} 0 & \text{if } \ell \leq R_{c1}, \\ \frac{\ell - R_{c1}}{(R_{c2} - R_{c1})} & \text{if } R_{c1} < \ell \leq R_{c2}, \\ 1 & \text{else.} \end{cases} \qquad [85]$$

A minimum size $R_{c1}$ is required for the cell to start proliferation, and the size is bounded from above by a maximum possible size of $R_{c2}$. Therefore,$R_{c1}$ matches $\sigma_h$ in our microscopic model.

We will compare the probability $p(t)$ that the cell has not proliferated at time $t > 0$ in the time frames $t_h$ to $t_\rho$, and in the regime $t > t_\rho$. As a starting point, we know how $p(t)$ behaves qualitatively in those two regimes from our analytical derivation, namely

$$p(t) \propto \begin{cases} \exp\left(-\frac{c_d(\sigma(t)-\sigma_h)^2}{2c_g}\right) & \text{if } t_h < t \leq t_\rho \\ \exp\left(-\frac{c_d(\sigma_\rho-\sigma_h)^2}{2c_g}\right) \cdot \exp\left(-(t-t_\rho)\cdot c_d(\sigma_\rho - \sigma_h)\right) & \text{if } t_\rho < t \end{cases} \qquad [86]$$

We now determine the evolution of $p_{\text{DPD}}(t)$ to determine $c_d$ such that the two distributions, from the simulations and from the microscopic model, match best. For simplicity, let us assume that $l_\rho \ll R_{c2}$ (which is the case in our simulations) and $R_{c2} - R_{c1} = N c_g \Delta t$.

Then we can calculate for the probability at $t = t_h + n\Delta t < t_\rho$, $n$ being the number of step after reaching $t_h$, the latter being the time at which $R_{c1}$ is reached. One gets

$$p_{\text{DPD}}(t_h + n\Delta t) = p_{\text{DPD}}(t_h) \prod_{i=1}^{n} \left(1 - P(R_{c1} + i\, c_g \Delta t)\right), \qquad [87]$$

where $P$ is defined by eq. (85) and equals one when $i = 0$. Follows,

$$\begin{aligned} p_{\text{DPD}}(t_h + n\Delta t) &= p_{\text{DPD}}(t_h) \prod_{i=0}^{n} \left(1 - P(R_{c1} + i c_g \Delta t)\right), \\ &= p_{\text{DPD}}(t_h) \prod_{i=0}^{n} \left(1 - \frac{i\, c_g \Delta t}{N c_g \Delta t}\right), \\ &= p_{\text{DPD}}(t_h) \prod_{i=0}^{n} \left(\frac{N-i}{N}\right), \\ &= p_{\text{DPD}}(t_h) N^{-n} \frac{N!}{(N-n-1)!}. \end{aligned}$$

Using the Stirling approximation leads to further simplifications,

$$\begin{aligned} p_{\text{DPD}}(t_h + n\Delta t) &\approx p_{\text{DPD}}(t_1) N^{-n} \frac{N!}{\sqrt{2\pi(N-n-1)}\left(\frac{N-n-1}{e}\right)^{N-n-1}}, \\ &= p_{\text{DPD}}(t_h) N^{-n} N! \frac{e^{N-n-1}}{\sqrt{2\pi(N-n-1)} e^{\ln(N-n-1)(N-n-1)}}, \\ &\approx p_{\text{DPD}}(t_h) \frac{N^{-n} N!}{\sqrt{2\pi N}} e^{N-n-1-\ln(N-n-1)(N-n-1)}, \end{aligned}$$

where the approximation $N - n - 1 \approx N$ has been used. Expanding the logarithm in the exponential function using Taylor expansion, one gets

$$\begin{aligned} p_{\text{DPD}}(t_h + n\Delta t) &\approx p_{\text{DPD}}(t_h) \frac{N!}{\sqrt{2\pi N}} N^{-n} e^{N-n-1-(\ln(N) - \frac{n+1}{N})(N-n-1)}, \\ &\approx p_{\text{DPD}}(t_h) \frac{N!}{\sqrt{2\pi N}} e^{-n\ln(N)+N-n-1-\left(\ln(N)(N-1) - n\ln(N) - (n+1) + \frac{(n+1)^2}{N}\right)}, \\ &\approx p_{\text{DPD}}(t_h) \frac{N!}{\sqrt{2\pi N}} e^{N-\left(\ln(N)(N-1)+\frac{(n+1)^2}{N}\right)}. \end{aligned}$$

In order to compare the simulations with eq. (86), we only consider factors containing $n$. Indeed, $n = (\sigma(t) - R_{c1})/c_g \Delta t$. Moreover, one has $N = (R_{c2} - R_{c1})/c_g \Delta t$ which leads to

$$\begin{aligned} p_{\mathrm{DPD}}(t_h + n\Delta t) &\propto e^{-\frac{(n+1)^2}{N}}, \\ &= \exp\left( - \frac{\left( \frac{\sigma(t) - R_{c1}}{c_g \Delta t} + 1 \right)^2}{\frac{R_{c2} - R_{c1}}{c_g \Delta t}} \right) \\ &= \exp\left( - \frac{(\sigma(t) - R_{c1} + c_g \Delta t)^2}{c_g \Delta t (R_{c2} - R_{c1})} \right) \\ &= \exp\left( - \frac{(\sigma(t + \Delta t) - R_{c1})^2}{c_g \Delta t (R_{c2} - R_{c1})} \right) \end{aligned}$$

Comparing this last formula with eq. (86), one gets the expression of the corresponding rate $c_d$

$$c_d = \frac{1}{(R_{c2} - R_{c1})\Delta t}. \qquad [88]$$

In simulation, the value of $R_{c1}$ being $R_{c1} = 0.75$ and the range of $R_{c2}$ being $R_{c2} \in [10^2, 2 \times 10^5]$, one can also write

$$c_d \approx \frac{1}{R_{c2} \Delta t}. \qquad [89]$$

**Table S2. Fitting parameters used for comparing eq. (2) and eq. (22) to the DPD simulations. Basis for the plots in fig 3c in the main manuscript.**

| Regime | $\mathbf{R_{c2}}$ **[s.u.]** | $\tau_d$ [s.u.] | $t_r$ [-] | $\sigma_r$ [-] | $\rho_h = \sigma_c^{-1}$ [s.u.] |
|---|---|---|---|---|---|
| gel-like | $1 \times 10^4$ | $0.25$ | 0.39 | 1.3529 | 0.68 |
| glass-like | $2 \times 10^5$ | $1.0$ | 2.51 | 1.8498 | 0.63 |

#### S8. Motivation of the Delayed Fisher Kolmogorov Formalism

In the main article, we propose an adapted continuous model for describing the macroscopic, i.e. tissue-wide, average dynamics of tissue. This model is based on the Fisher-Kolmogorov (FK) differential equation, which describe the evolution of the local cell density $\rho$ as a function of time and space as:

$$\partial_t \rho = D\Delta\rho + \beta(\rho_h - \rho)\rho \qquad [90]$$

with the two main parameters $D$ and $\beta$. Here $D$ is the diffusion coefficient describing the tissue expansion, which is driven by an internal pressure build up due to increasing cell density. The associated term is referred to as the *diffusive term*. The parameter $\beta$ corresponds to the growth coefficient controlling the rate at which cell density converges to its equilibrium density $\rho_h$, which is an implicit parameter of the model. The associated term is referred to the *proliferation* or *growth* term.

In the classical FK formulation (eq. (90)), cell division is instantaneous, as the density gained from cell division in the growth term is instantly added to the total density, which can in turn immediately contribute to the proliferation of the formula again. This observation does neither agree well with our observations in experiments, where it is clear that the process of DNA duplication and formation of a new cell membrane does take time, nor with our predictions by the cell-level model, where the finite time $\tau_d$ is essential to declare whether a cell is dividing or not. Moreover, one can scale the ratio of proliferating cells $\mathcal{R}$ for the FK equation. The second term on the r.h.s. of eq. (90) denotes the cells dividing during an infinitesimal time step $\delta t$ in a given infinitesimal volume $\delta V$, i.e.

$$\delta N = \beta(\rho_h - \rho)\rho\, \delta t\, \delta V. \qquad [91]$$

The amount of cell in the same volume is $N = \rho\, \delta V$, leading to

$$\mathcal{R} \approx \delta N/N = \beta(\rho_h - \rho)\delta t. \qquad [92]$$

This expression does not account for the change of curvature around $\rho_h$ observed experimentally and therefore fails to account for mechanosensitive properties.

In terms of differential equations, these observations have several consequences. Firstly, the effect of the growth term has to be delayed, leaving us with a delayed differential equation. Secondly, to keep track of the portion of cells currently dividing and the subpopulation still growing and available for division, we need to distinguish these two populations with different states. Finally, to capture mechanosensing, one needs to add an exponent to the division term of the FK equation. This new exponent can then be seen as a scaling-behavior, for how the rate of proliferation speeds up or slows down as the local density approaches $\rho_h$. Consequently, this exponent will allow to control the curvature of $\mathcal{R}(\rho)$ close to $\rho_h$. Overall, we have the following system of coupled delayed Fisher-Kolmogorov (DFK) differential equations:

$$\text{Growing}: \quad \partial_t \rho_g(\vec{r},t) = D\Delta\rho_g(\vec{r},t) \underbrace{-\beta\left(1-\frac{\rho(\vec{r},t)}{\rho_0}\right)^c \cdot \rho_g(\vec{r},\tau)}_{\text{Entering division}} \underbrace{+2\cdot\beta\left(1-\frac{\rho(\vec{r},\tau)}{\rho_0}\right)^c \cdot \rho_g(\vec{r},\tau)\bigg|_{\tau=t-\tau_s}}_{\text{Leaving division}} \qquad [93]$$

$$\text{Dividing}: \quad \partial_t \rho_d(\vec{r},t) = \underbrace{\beta\left(1-\frac{\rho(\vec{r},t)}{\rho_0}\right)^c \cdot \rho_g(\vec{r},t)}_{\text{Entering division}} - \underbrace{\beta\left(1-\frac{\rho(\vec{r},\tau)}{\rho_0}\right)^c \cdot \rho_g(\vec{r},\tau)\bigg|_{\tau=t-\tau_s}}_{\text{Leaving division}} \qquad [94]$$

#### S9. Details of the DDE solver and the numerical solution to the DFK

To obtain the numerical solution of the DFK equations (eqs. (93) and (94)), we implemented a custom python script to compute the time evolution of the density distributions $\rho_{g,d}$ for a given set of parameters ($\rho_h$, $D$, $\tau_d$, $\beta$, $c$) by solving a radially symmetric tissue with $\rho(\vec{r}, t) = \rho(r, t)$ in two dimensions (determines the appropriate Laplace operator).

$$\partial_t \rho_g(r,t) = D\left(\partial_r^2 + \frac{1}{r}\partial_r\right)\rho_g(r,t) - \beta\left(1 - \frac{\rho(r,t)}{\rho_h}\right)^c \rho_g(r,\tau) + 2\beta\left(1 - \frac{\rho(r,\tau)}{\rho_h}\right)^c \rho_g(r,\tau)\Big|_{\tau=t-\tau_d}, \quad [95]$$

$$\partial_t \rho_d(r,t) = \beta\left(1 - \frac{\rho(r,t)}{\rho_h}\right)^c \rho_g(r,t) - \beta\left(1 - \frac{\rho(r,\tau)}{\rho_h}\right)^c \rho_g(r,\tau)\Big|_{\tau=t-\tau_d}. \quad [96]$$

Using the polar representation of the Laplace operator together with $\partial_\phi \rho_{g,d} = 0$, i.e. a homogeneous tissue along the azimuthal direction, the radial components of the DFK equations are given by eqs. (95) and (96). The equations are integrated on an interval $r \in [0, R]$ with reflecting boundary conditions at the origin, i.e. $\partial_r \rho_{g,d}(r = 0, t) = 0, \forall t$, and absorbing boundary conditions at the outer boundary, i.e. $\rho_{g,d}(r = R, t) = 0$. As initial conditions, we assume a static density distribution with no proliferation, namely

$$\rho_g(r, t \le 0) = \begin{cases} \rho_{\text{init}} & \text{if } r \le r_{\text{init}}, \\ 0 & \text{else,} \end{cases} \quad [97]$$

$$\rho_d(r, t \le 0) = 0. \quad [98]$$

Spatial discretization is performed with a discrete step of $\Delta r$ leading to discrete positions $r_i = i\,\Delta r$. The numerical derivatives are calculated by central differences, i.e. $\partial_r f(r_i) = (f(r_{i+1}) - f(r_{i-1}))/(2\Delta r)$, while the Laplace operator is implemented as the usual central stencil, i.e. $\Delta_r f(r_i) = (f(r_{i+1}) - 2f(r_i) + f(r_{i-1}))/(\Delta r)^2$.

The numerical solution of the time evolution of $\rho_{g,s}$ is obtained via a custom SciPy-based python module *ddesolver* (https://github.com/Reshief/ddesolver), which is a modification of the python module *ddeint* (https://github.com/Zulko/ddeint) with higher numerical accuracy thanks to Runge-Kutta 4/5 integration steps and time-step rescaling. To account for the delayed term of the DFK, a history of states for simulated times $t \ge 0$ is kept. This history is looked into during integration, and linear interpolation is used between subsequent simulated time steps. Such history repositories are implemented for both densities $\rho_g$ and $\rho_d$ independently. The simulation is then carried out for a total time $T > 0$ with a time discretization step of a maximum of $\Delta t$ in arbitrary units.

The values of simulation parameters are summarized into table S3. The data for $R(\rho)$ is obtained by time-averaging and binning of the ratios $\rho_s/\rho$ as a function of $\rho$ at all positions with $\rho_0 \ge \rho \ge 0.01$ with 40 equidistant $\rho$-bins on the interval $[0, \rho_0]$.

**Table S3. Values of parameters chosen for numerical integration of the proposed DFK system of equations presented in fig. S9 and fig. S8. If multiple values are listed, all combinations including each of these values with all possible combinations of listed values for other parameters have been simulated. For fig. 3d in the main manuscript, the parameters except for $c$ were fixed to $D = 1$, $\beta = 1$, $\tau_d = 1$ and $c$ was varied between $0.25$ and $2$ in increments of $0.25$.**

| Parameter name | Values |
|---|---|
| $D$ | $0.25, 0.5, 1, 2, 3$ |
| $b$ | $0.25, 0.5, 0.75, 1.0, 1.5, 2.0, 2.5, 3.0$ |
| $c$ | $0.25, 0.5, 0.75, 1.0, 1.25, 1.5, 1.75, 2.0$ |
| $\tau_s$ | $0.25, 0.5, 1, 2, 3, 4, 5$ |
| $R$ | $100$ |
| $r_{init}$ | $5$ |
| $\rho_{init}$ | $1.0$ |
| $\rho_0$ | $1.0$ |
| $\Delta t$ | $0.01$ |
| $t$ | $100$ |
| $\Delta r$ | $0.25$ |

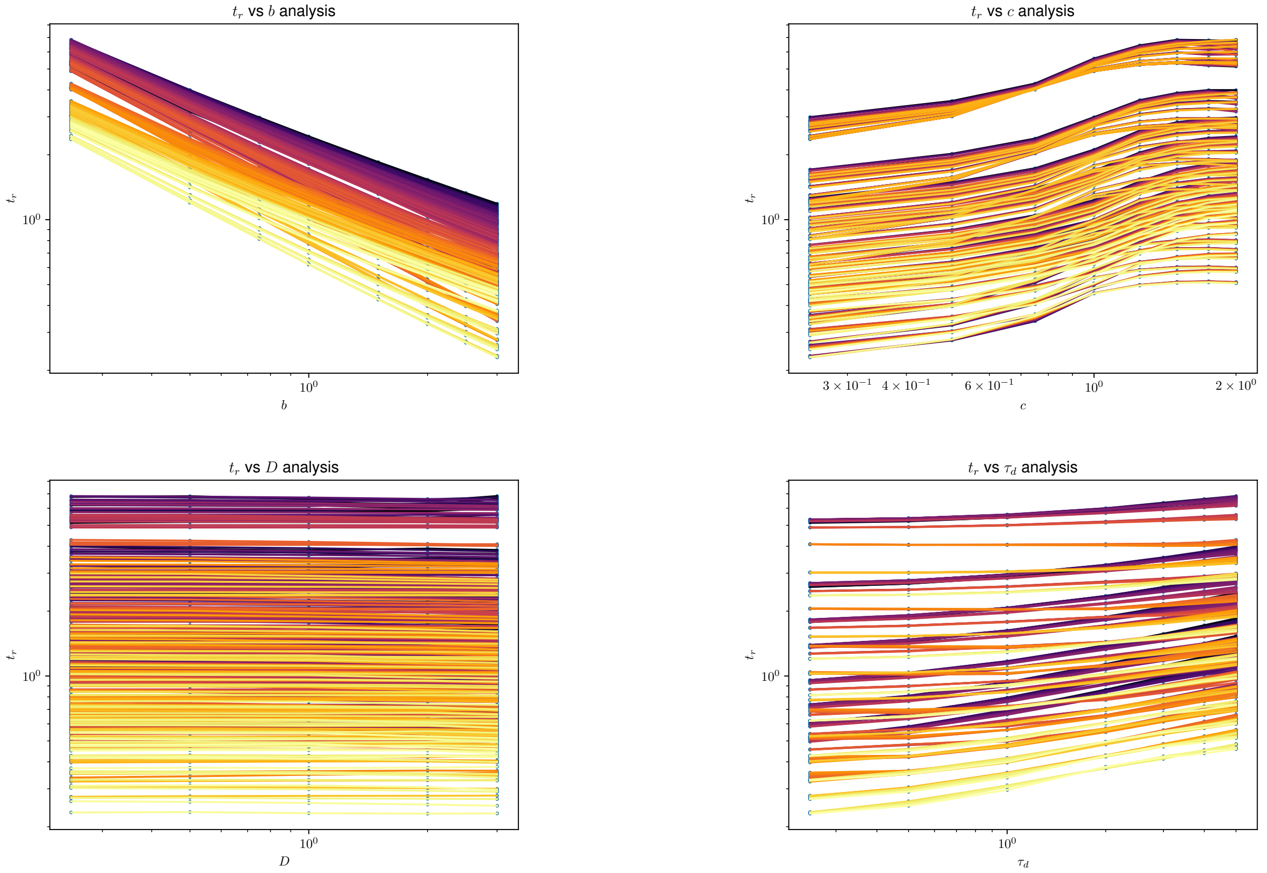

**Fig. S8. Illustrations of the link between the DFK parameters and the cell-level parameter $t_r$.** Plots of the results for $t_r$ when fitting the cell-level model to $\mathcal{R}$ profiles of the DFK simulations. We generally observe a $t_r \propto 1/\beta$ relation in the data as well as a slight link to $c$ but not much of a connection to the parameter $D$ of the DFK. The slight link to $c$ seems to be related to the effect that $\sigma_r$ has on the average $\tau_g$ at low densities, which makes $\sigma_r$ and $c$ not fully disconnected from the time scale $t_r$. The dependence on $\tau_d$ is a consequence of the fit of $\mathcal{R}$ in terms of $t_r/\tau_d$ and numeric instabilities at high values of $\tau_d$.
Lines in each graph indicate constant sets of all other parameters with only the parameter on the $x$-axis being varied. Different lines are assigned different colors for better visual differentiation.

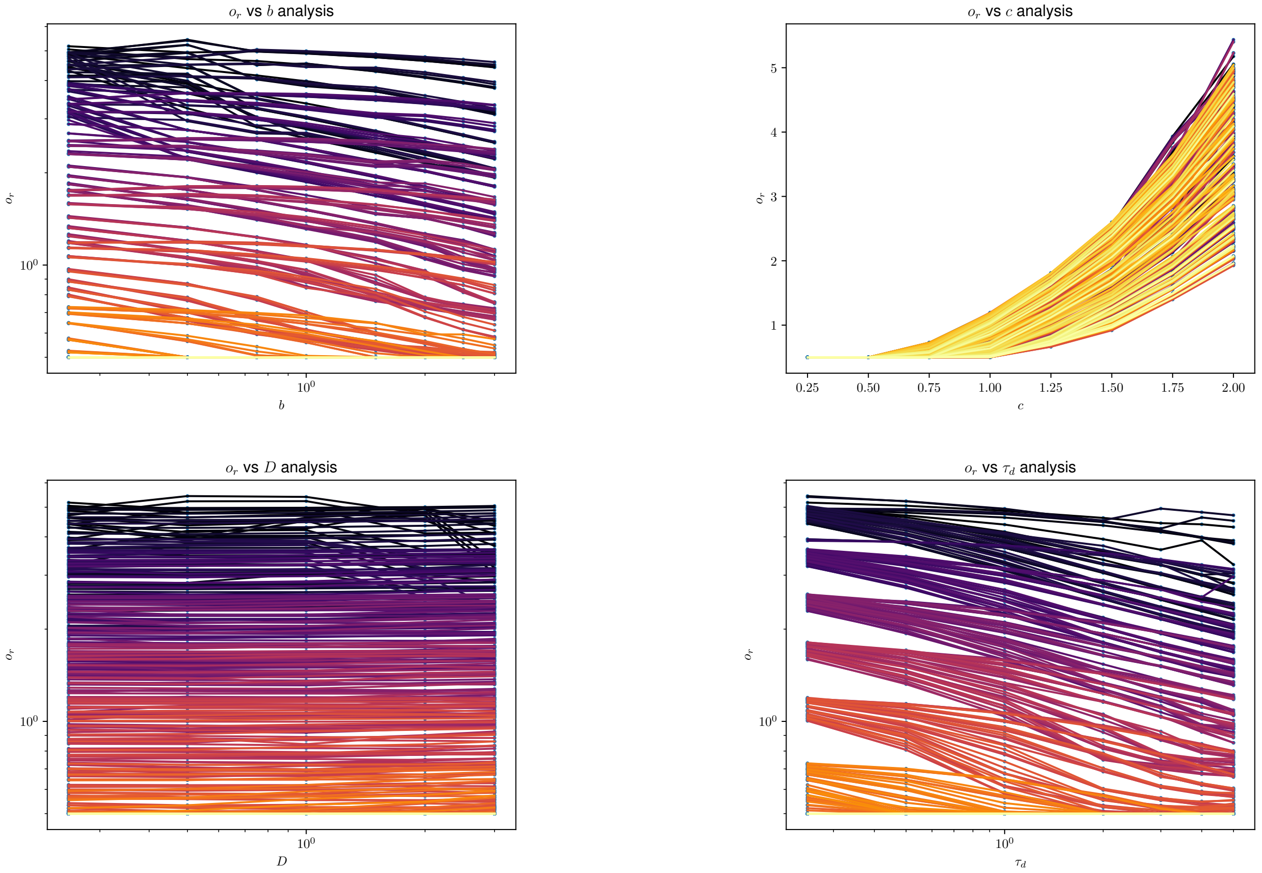

**Fig. S9. Illustrations of the link between the DFK parameters and the cell-level parameter $\sigma_r$.** Plots of the results for $\sigma_r$ when fitting the cell-level model to $\mathcal{R}$ profiles of the DFK simulations. The only conclusive connection we observe is that between $\sigma_r$ and $c$ (and implicitly also a linear scaling with $\sigma_h$ as the reference size scale) with no consistent impact of other parameters of the DFK on $\sigma_r$. This agrees well with our understanding, that $\sigma_r$ and $c$ are the parameters controlling the curvature of $\mathcal{R}$ in their respective models.
Lines in each graph indicate constant sets of all other parameters with only the parameter on the $x$-axis being varied. Different lines are assigned different colors for better visual differentiation.

#### S10. Behavior of $\tau_g$ and $\mathcal{R}$ around $\rho_h$

Equation 22 can be developed as a series around the homeostatic density $\rho_h$. Indeed, assuming that the background density $\rho$ is such as $\rho_h - \delta\rho$, first of all, we obtain a linear link between $\Delta\sigma$ and $\delta\rho$ (where we use $b = 0.1$ for better readability)

$$\Delta\sigma = \frac{2 - (\rho\sigma_h)^{0.1}}{\rho} - \sigma_h \quad [99]$$

$$= \frac{2 - ((\rho_h - \delta\rho)\sigma_h)^b}{\rho_h - \delta\rho} - \sigma_h \quad [100]$$

$$\approx \sigma_h^2(b+1)\delta\rho - \frac{\sigma_h^3(b^2 - 3b - 2)}{2}\delta\rho^2 + \mathcal{O}\left(\delta\rho^3\right) \quad [101]$$

and consequently eq. (22) can be written as

$$\langle\tau_g\rangle = t_r\left[\frac{\sigma_h}{2\sigma_r} + \frac{\sigma_r}{\Delta\sigma}\right] + \mathcal{O}(\Delta\sigma^3) \quad [102]$$

$$[103]$$

which indicates a hyperbolic divergence of the growth time around the homeostatic density. Continuing with $\mathcal{R}$, we arrive at:

$$\mathcal{R} = \frac{\tau_d}{\tau_d + \tau_g} \quad [104]$$

$$= \frac{1}{1 + \frac{t_r}{\tau_d}\cdot\left[\frac{\sigma_h}{2\sigma_r} + \sqrt{\pi/8}\times\mathrm{erf}\left(\frac{\Delta\sigma}{\sqrt{2}\sigma_r}\right) + \frac{\sigma_r}{\Delta\sigma}\exp\left(-\frac{(\Delta\sigma)^2}{2\sigma_r^2}\right)\right]} \quad [105]$$

$$\approx \frac{\Delta\sigma}{\sigma_r}\cdot\frac{\tau_d}{t_r} - \frac{\Delta\sigma^2\sigma_h}{2\sigma_r^3}\cdot\left(2\frac{\sigma_r}{\sigma_h} + \frac{t_r}{\tau_d}\right)\cdot\frac{\tau_d^2}{t_r^2} + \mathcal{O}\left(\Delta\sigma^3\right) \quad [106]$$

Now, plugging in the series expansion of $\Delta\sigma$ into that of $\mathcal{R}$, we arrive at:

$$\mathcal{R}(\rho_h - \delta\rho) \approx \frac{\sigma_h^2(b+1)\delta\rho - \frac{\sigma_h^3(b^2-3b-2)}{2}\delta\rho^2}{\sigma_r}\cdot\frac{\tau_d}{t_r} - \frac{(b+1)^2\delta\rho^2\sigma_h^5}{2\sigma_r^3}\cdot\left(2\frac{\sigma_r}{\sigma_h} + \frac{t_r}{\tau_d}\right)\cdot\frac{\tau_d^2}{t_r^2} + \mathcal{O}\left(\delta\rho^3\right) \quad [107]$$

$$= \delta\rho\cdot\frac{\sigma_h^2(b+1)}{\sigma_r}\cdot\frac{\tau_d}{t_r} + \delta\rho^2\cdot\frac{\sigma_h^3}{2\sigma_r}\cdot\frac{\tau_d}{t_r}\left[\frac{(2+3b-b^2)}{2} - \frac{\sigma_h^2(b+1)^2}{\sigma_r^2}\cdot\left(2\frac{\sigma_r}{\sigma_h} + \frac{t_r}{\tau_d}\right)\cdot\frac{\tau_d}{t_r}\right] + \mathcal{O}\left(\delta\rho^3\right) \quad [108]$$

As a result, one sees that the behavior of $\mathcal{R}$ around $\rho_h$ is polynomial, and has a curvature which depends on the sign of the term in brackets.
